# Heat Stress Regulates the Expression of *TPK1* Gene at Transcriptional and Post-Transcriptional Levels in *Saccharomyces cerevisiae*

**DOI:** 10.1101/2021.10.07.463258

**Authors:** Luciana Cañonero, Constanza Pautasso, Fiorella Galello, Lorena Sigaut, Lia Pietrasanta, Arroyo Javier, Mariana Bermúdez Moretti, Paula Portela, Silvia Rossi

## Abstract

In *Saccharomyces cerevisiae,* cAMP regulates a number of different cellular processes, such as cell growth, metabolism, stress resistance and gene transcription. The intracellular target for this second messenger in yeast cells is the cAMP-dependent protein kinase (PKA). The way in which a broad specificity protein kinase mediates one right physiological response after cAMP increase indicates that specificity is highly regulated in the cAMP / PKA system. Here we address the mechanism through which cAMP-PKA signalling mediates its response to heat shock thermotolerance in *Saccharomyces cerevisiae*. Yeast PKA is a tetrameric holoenzyme composed of a regulatory (Bcy1) subunit dimer and two catalytic subunits (Tpk1, Tpk2 and Tpk3). PKA subunits are differentially expressed under certain stress conditions. In the present study we show that, although the mRNA levels of *TPK1* are upregulated upon heat shock at 37°C, no change is detected in Tpk1 protein levels. The half-life of *TPK1* mRNA increases and this mRNA condensates in cytoplasmic foci upon thermal stress. The resistance of *TPK1* mRNA foci to cycloheximide-induced disassembly, together with the polysome profiling analysis suggest that *TPK1* mRNA is impaired for entry into translation. *TPK1* mRNA foci and *TPK1* expression were also evaluated during thermotolerance. The crosstalk of cAMP-PKA pathway and cell wall integrity (CWI) signalling was also studied. Wsc3 sensor and other components of the CWI pathway are necessary for the upregulation of *TPK1* mRNA upon heat shock conditions. The assembly in cytoplasmic foci upon thermal stress shows to be dependent of Wsc3. Finally, evidence of an increase in the abundance of Tpk1 in the PKA holoenzyme in response to heat shock is presented, suggesting that a recurrent stress enhanced the fitness for the coming favorable conditions The results indicate the existence of a mechanism that exclusively regulates Tpk1 subunit expression and therefore contributing to the specificity of cAMP-PKA.

**SUMMARY STATEMENT:** PKA subunits are differentially expressed under heat-shock conditions. The mRNA of the TPK1 subunit is upregulated upon heat-shock at 37°C and thermotolerance, the half-life increases upon heat-stress and also this transcript condensates in cytoplasmic foci upon thermal stress and thermotolerance. The resistance to cycloheximide treatment of TPK1 mRNA foci together with the analysis by polysome profiling suggest that TPK1 mRNA is impaired for entry into translation upon thermal stress. An increase in Tpk1 protein and PKA activity was detected after the heat stress treatments.

Cell Wall Integrity pathway, through Wsc3 sensor, is involved in TPK1 expression. Heat-stress regulates TPK1 expression through this pathway from an intermediate step of the cascade and independently of the upstream elements of the CWI pathway activation. These results demonstrate a new crosstalk between the two signalling pathways.

The increment in Tpk1-dependent PKA activity during cell adaptation to heat stress might contribute to the overall cellular fitness when more favorable environmental conditions are restored.

The results indicate the existence of a mechanism that exclusively regulates TPK1 subunit expression and therefore contributing to the specificity of cAMP-PKA.

## INTRODUCTION

The yeast *Saccharomyces cerevisiae* has developed sensing systems and complex signalling networks to efficiently respond to sudden and frequent variations in temperature, osmolarity, environmental acidity, presence of toxins and long periods of nutritional starvation (Hohmann and Mager, 2003). These adaptive circuits control the expression of numerous genes that are involved in cell division, metabolic pathways, stress resistance and cell differentiation (Berry and Gasch, 2008; De Virgilio and Loewith, 2006; Zaman et al., 2008).

The inhibition of protein translation plays a key role in the stress adaptation process, saving energy in this consuming biosynthetic pathway. At the same time, the upregulation of the translational initiation of specific mRNAs leads to the induction of certain proteins, allowing adaptation (Crawford and Pavitt, 2019; Simpson and Ashe, 2012).

The cAMP-PKA signalling pathway is involved in the precise coordination of cellular responses to various stimuli (Conrad et al., 2014; Thevelein et al., 2008). How signalling specificity is attained when different stimuli trigger the endogenous production of the same second messenger, cAMP, is not entirely understood. *S. cerevisiae* PKA is composed of two regulatory subunits encoded by a single gene *BCY1,* and two catalytic subunits encoded by the *TPK1*, *TPK2* and *TPK3* genes. Considering the pleiotropic role of PKA in unicellular organisms, and the great diversity of substrates at different subcellular sites, regulatory mechanisms must exist to ensure the phosphorylation of specific substrates under certain stimulus. The different catalytic subunits (Tpks) seem to be functionally redundant for cell viability although several specific functions for each isoform have been described (Palomino et al., 2006; Pan and Heitman, 2002; Robertson and Fink, 1998; Robertson et al., 2000). As was previously shown, cAMP-PKA signal transduction specificity is not only accomplished by the recognition of substrate consensus sequences for each catalytic PKA subunit (Galello et al., 2010; Mok et al., 2010). The specificity of the response is achieved by several levels of control which act coordinately. One of these control levels is the restriction of the localization of each catalytic subunit to subcellular compartments defined by the interaction of the regulatory subunit with anchoring proteins. In mammals, A-Kinase Anchoring Proteins (AKAPs) have been described as spatiotemporal modulators of cAMP-dependent protein kinase activity (Calejo and Taskén, 2015; Dema et al., 2015; Pidoux and Taskén, 2010; Skroblin et al., 2010). However, in *S. cerevisiae*, the existence of AKAPs has been demonstrated. The localisation of Bcy1 is dynamic and responsive to environmental and nutritional conditions (Griffioen et al., 2000; Tudisca et al., 2010). Several N-terminal Bcy1 dependent interacting proteins have been described in *S. cerevisiae*. However, the structural determinants necessary for interaction are different from those of mammalian AKAPs (Galello et al., 2014). In mammals there are also evidence that indicates the targeting of the catalytic subunits to several proteins in the cytosol and the nucleus (Søberg and Skålhegg, 2018).The differential localization of each Tpk isoform in yeast was demonstrated in fermentative, respiratory, and stationary phases of growth, as well as under several stress conditions (Portela and Rossi, 2020). It has been shown that Tpk2 and Tpk3 isoforms are differently associated with processing bodies (P-bodies) and stress granules (SGs) during stationary phase, glucose starvation, heat stress and hyper osmotic stress (Tudisca et al., 2010).

The regulation of the expression of each PKA subunit is another important aspect regarding signal transduction specificity in which we have also been working on. Some antecedents, in mammals, indicate that cAMP modulates the expression of PKA subunits (Houge et al., 1990; Hougel, 1992; Knutsen et al., 1991; Taskén et al., 1991). In yeast, our antecedents indicate that all PKA subunits share an autoregulatory and inhibitory mechanism mediated by PKA activity (Pautasso and Rossi, 2014). However, each PKA subunit also presents a distinctive transcription during the growth phase, the growth in non-fermentable or fermentable conditions and upon heat shock or saline stress (Galello et al., 2017; Pautasso and Rossi, 2014). *TPK1* is the only PKA subunit that is transcriptionally upregulated during heat shock and saline stress (Pautasso and Rossi, 2014). Moreover, *TPK1* promoter, but not *TPK2* and *TPK3*, presents positioned nucleosomes that are evicted upon heat stress (Reca et al., 2020). We have previously described a complex network of specific expression regulators for each PKA subunit with a high-throughput screening (Pautasso et al., 2016). These results expand our understanding of the importance of transcriptional regulation of PKA subunits in controlling the specificity of cAMP-PKA signalling. Nevertheless, the increase in *TPK1* transcription during stress appears paradoxical considering that higher PKA activity in yeast cells leads to lower stress resistance (Thevelein et al., 2000). This would suggest the existence of an intriguing mechanism that regulates the expression of PKA subunits. Several mechanisms can allow organisms to be ready for recurring stressors. One of them is the anticipatory response or cellular memory, through which a current environment acts as a signal or input, resulting in adaptation to future challenges (Jiang et al., 2020).

Different environmental stresses may trigger some overlapping cellular responses involving HOG (High Osmolarity Glycerol), CWI (Cell Wall Integrity), HSR (Heat Shock Response) or ESR (Environmental Stress Response) pathways (Engelberg et al., 2014; Morano et al., 2012; Santiago et al., 2020). Exposure to high temperatures leads to the activation of the CWI, the ESR and the HSR pathways. HSR can be considered a subset of the ESR and is governed by the action of primarily two transcription factors, Hsf1 and Msn2/4. *TPK1* shows to be dependent on Msn2/4 but not on Hsf1 upon heat-shock (Pautasso and Rossi, 2014). Thermal stress not only affects internal cellular processes but also impacts the cell surface. It has been shown that CWI, a complex signalling pathway, links transmembrane proteins with a kinase cascade that ends in the activation of effector transcription factors. The CWI pathway is activated in response to perturbations in the cell ultrastructure, including compounds that interfere with cell wall synthesis and changes in pH and temperature (Engelberg et al., 2014). Several CWI sensors have been described, namely Wsc1–3, Mid2 and Mtl1 (Kock et al., 2015; Rodicio and Heinisch, 2010). These sensors recruit Rom1/ Rom2 (guanine nucleotide exchange factors) which activates the small G-protein Rho1. The Rho1-GTP then stimulates the downstream effector Pkc1, which regulates the MAPK cascade (Bck1 and Mkk1/2). Finally, the kinases Mkk1/2 activate the MAPK Slt2 (or Mpk1). Slt2 controls two transcription factors, Rlm1 and Swi4/6, which trigger the transcription of genes involved in cell wall biogenesis (Levin, 2011; Verghese et al., 2012). Temperature increases activates the Slt2 kinase to restore cell membrane fluidity. In the absence of the sensors Mid2 or Wsc1-4, the HSR is activated, but cells are heat shock sensitive, autolytic and the CWI transcription factor Rlm1 is not activated.

During stress, reprogramming of gene expression occurs, which involves the global inhibition of translational initiation and the large-scale induction of stress-responsive mRNAs (Causton et al., 2001; Crawford and Pavitt, 2019; Gasch et al., 2000). In addition, coordinated regulation of mRNA abundance, translation, localization and turnover rates of individual mRNAs have been described during several stresses such as heat shock (Crawford and Pavitt, 2019; Halbeisen and Gerber, 2009; Preiss et al., 2003). Therefore, both transcriptional and translational regulations contribute to stress-induced changes in gene expression, albeit the exact effect of each process and the equilibrium between them depend on the type of stress.

In this post-transcriptional control, the specific localization and compartmentalization of mRNAs within the cytoplasm plays an important role (Gehring et al., 2017; Pizzinga and Ashe, 2014). Particular sets of proteins and translationally repressed mRNAs colocalize in cytoplasmic discrete ribonucleoprotein (RNP) foci that differ in composition according to various external and internal signals. P-bodies and SGs contain mRNAs and specific but overlapping proteins as constituents (Corbet and Parker, 2020). P-bodies are defined as sites of mRNA storage and/or decay. SGs represent a reservoir of translational inactive mRNAs, translation factors and associated proteins (Mittag and Parker, 2018). Other granules have been described in yeast associated with different biological activities, like the foci of several proteins involved in intermediary metabolism and stress response and actin bodies in quiescent cells (Narayanaswamy et al., 2009; Sagot et al., 2006). In this study, we analyse the regulation of *TPK1* expression during thermal stress adaptation. When cells grown at 25°C are subjected to heat shock treatment at 37°C for 60 minutes, the *TPK1* mRNA is upregulated while the protein levels show no change. However, when yeast cells are subjected to a scheme of temperature shifts (heat shock at 37°C (1) - recovery period at 25°C (1) - heat shock 37°C (2) - recovery period at 25°C (2)) to induce heat shock memory, an upregulation of Tpk1 protein expression can be detected in both recovery phases at 25°C. Upon incubation at 37°C, the half-life of *TPK1* mRNA increases and this mRNA is detected in cytoplasmic foci. These foci are disassembled when stress is removed but not after cycloheximide treatment. This suggests that these mRNAs are impaired for entry into translation. It was also demonstrated a crosstalk between cAMP-PKA pathway and CWI signalling in the regulation of *TPK1* expression. The upregulation of the *TPK1* during heat shock is dependent on the CWI pathway and the sensor Wsc3. Wsc3 is also necessary for the induction of *TPK1* mRNA cytoplasmic foci during heat shock. Finally, the proportion of the Tpk1 isoform in the holoenzyme is significantly higher during the second 25°C recovery period than at any of the other conditions tested. These results indicate a particular mechanism of Tpk1 subunit expression during thermal stress memory that contributes to holoenzyme conformation and thus to cAMP-PKA specificity.

## RESULTS

### *TPK1* mRNA decay kinetics during heat shock

We have previously demonstrated that the *TPK1* promoter, but not the other PKA subunit promoters, is activated upon heat shock and osmostress (Pautasso and Rossi, 2014; Reca et al., 2020). We also demonstrate that, during exponential growth, the *BCY1* promoter has the lowest activity of the four PKA subunit promoters and does not respond to thermal stress (Pautasso and Rossi, 2014). This discrepancy between the promoter activity and the transcript abundance could be the consequence of differences in mRNA stability. With this background in mind, we evaluated the mRNA stability and the protein levels of *TPK1* and *BCY1.* As none of the *TPK2*, *TPK3* or *BCY1* promoter activities or their mRNA levels significantly change after heat shock, we decided to assess only *BCY1* mRNA half-life and Bcy1 protein levels to compare with *TPK1* mRNA and protein levels (Pautasso and Rossi, 2014). Figure 1 shows the protein and mRNA levels of *TPK1* and *BCY1* upon heat shock stress. A three-fold increase in *TPK1* mRNA levels was determined upon stress while no change in Tpk1 protein levels was detected (Fig. 1A). On the other hand, neither Bcy1 protein nor *BCY1* mRNA levels showed any difference in response to thermal stress (Fig. 1C). *TPK1* and *BCY1* mRNAs decay rates were assessed in unstressed exponentially growing cells using the transcriptional inhibitor 1,10-phenanthroline. At 25°C, *BCY1* mRNA decayed with a t1/2 = 19.5 min, whereas *TPK1* mRNA decayed with a t1/2 = 4.6 min (Fig. 1B, D and E). This result indicated that *BCY1* mRNA is more stable than *TPK1* mRNA. To test whether the decay of these mRNAs is modified by heat stress, exponentially growing cells were incubated with the transcriptional inhibitor after the thermal treatment. Both *TPK1* and *BCY1* mRNAs were more stable upon stress than at 25°C, being both t1/2 values > 60 min (Fig. 1B, D and E).

**Figure 1.**
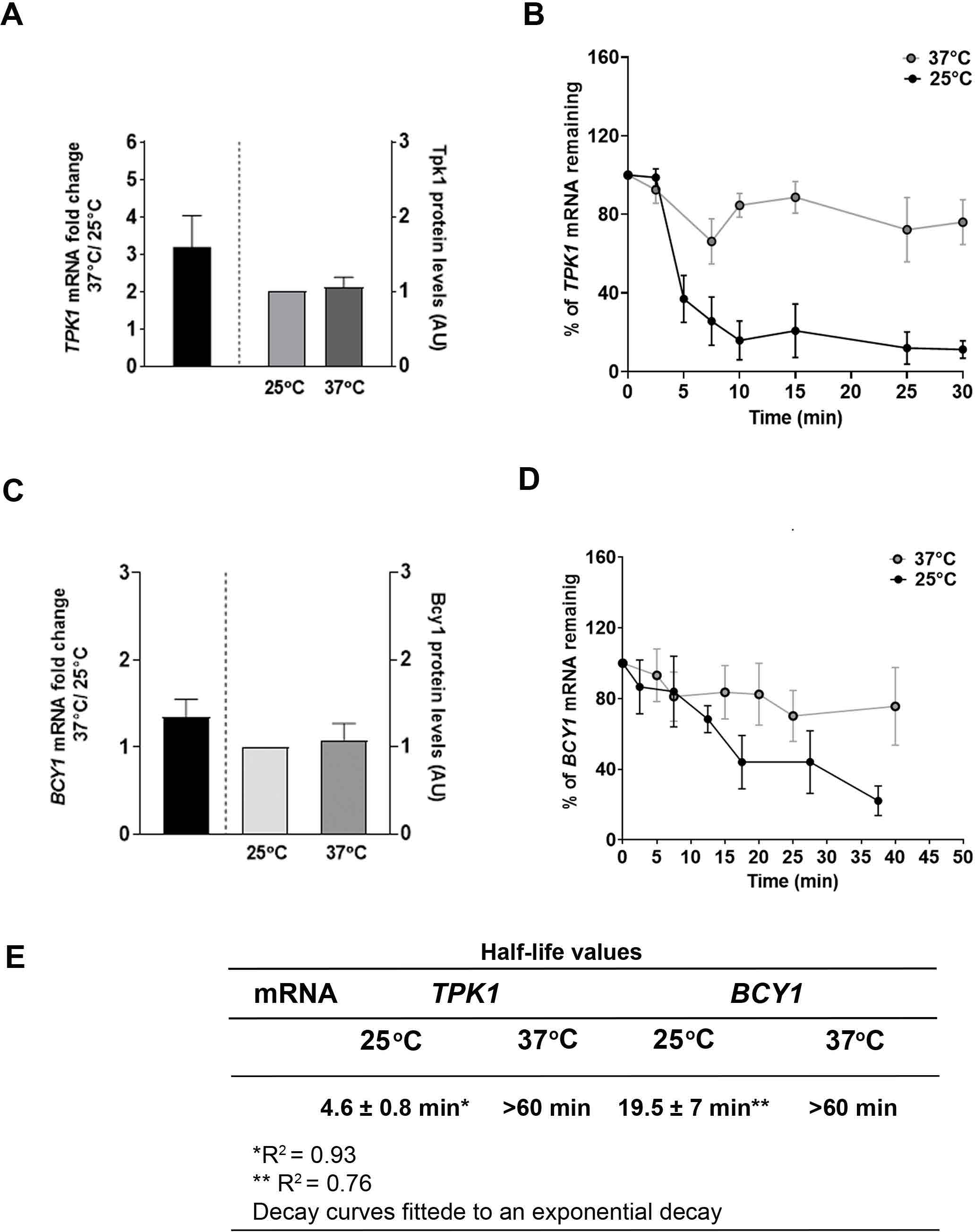
*TPK1* mRNA half-life but not Tpk1 protein is increased during heat shock. A) *TPK1* mRNA levels were assessed by RT-qPCR, normalized to *TUB1* mRNA and expressed as fold change 37°C/25°C (left panel). Protein extracts were analysed by Western blot with Tpk1 antibody, quantified and expressed as AU (arbitrary units) (right panel). B) Cells were incubated or not at 37°C for 60 min and samples were harvested at the indicated time points following transcriptional arrest by 1,10-phenanthroline. *TPK1* mRNA levels were meassured at the indicated time points after the addition of the drug using northern analysis. Band intensities were quantified by PhosphorImager. 28S and 18S rRNA were used as loading controls. The intensity at time 0 (before adding the drug) was defined as 100%, and the intensities at the other time points were calculated relative to time 0. The decay curves of each mRNA were fitted to an exponential decay model. Error bars represent s.e.m. of three assays. C) qRT-PCR was performed to measure *BCY1* mRNA and Western blot to quantify Bcy1 using Bcy1 antibody. D) Idem B, decay kinetics was determined by monitoring *BCY1* mRNA levels. E) Table summarising the t1/2 values for *TPK1* and *BCY1* mRNAs at 25ᵒC or 37ᵒC temperature.

As already mentioned in the introduction, organisms are continually challenged by changing environments and develop physiological adaptations to deal with such changes. Several mechanisms as cellular memory allow organisms to be ready for recurrent stressors (Jiang et al., 2020). To further understand the molecular process that mediates the regulation of *TPK1* expression, gene expression during thermotolerance was evaluated. To this aim we exposed the cells to a scheme of two consecutive heat shocks (60 min the first one and 30 min the second) each one followed by a period of recovery at 25°C (Fig. 2A). Then, protein and mRNA levels of *TPK1* and *BCY1* were assessed. As shown in Figure 2B, the Tpk1 protein level detected after each heat shock significantly increased during both recovery periods. It was also observed that during the first 25°C period, the level of Tpk1 protein was substantially lower than the level detected during the second one. In contrast, the levels of *TPK1* mRNA presented an inverted pattern, with a significant increase at 37°C (Fig. 2C, left panel). Bcy1 protein and *BCY1* mRNA levels remained unchanged throughout the entire experiment (Fig. 2B and C, right panels).

**Figure 2.**
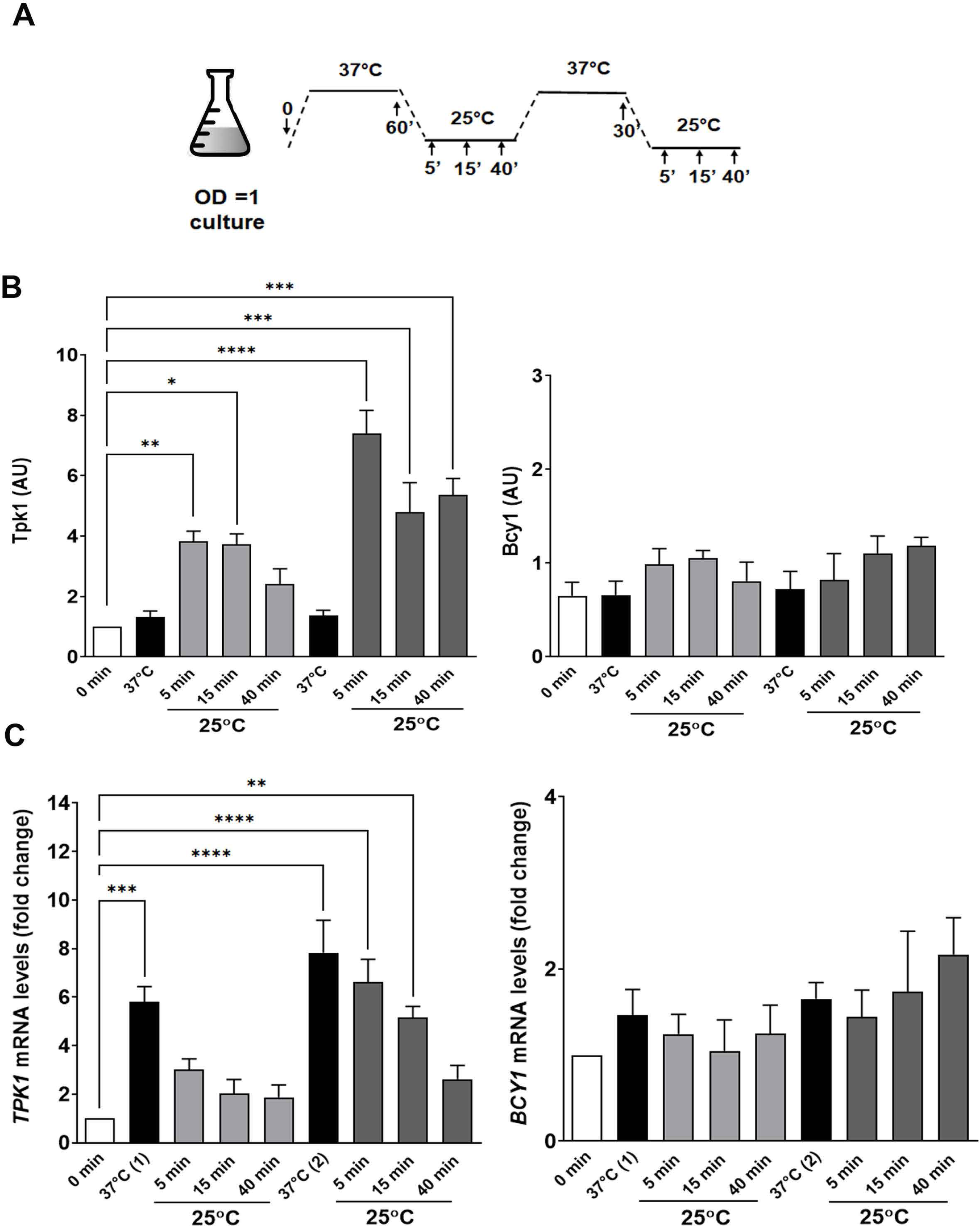
*TPK1* mRNA and Tpk1 protein have a particular expression pattern during heat shock adaptation. A) Scheme of temperature shifts applied. Cells were exposed to a two thermal stress at 37°C, the first one for 60 min and the second for 30 min, each one followed by a 25°C incubation. Samples were harvested at different times during each incubation. B) Western blot using anti-Tpk1 or anti-Bcy1 antibodies (right and left panels, respectively). C) *TPK1* mRNA and *BCY1* mRNA levels were measured by qRT-PCR. Results are expressed as mean ± s.e.m. from at least three independent experiments. Statistical analysis corresponds to two-way ANOVA, *p<0.05; **p<0.01; ***p<0.001; ****p<0.0001.

### *TPK1* mRNA associates with granules upon heat stress

It is known that higher PKA activity leads to lower stress resistance. However, our antecedents and the above results indicate that *TPK1* mRNA is upregulated and stabilized upon stress. Thus, we considered that *TPK1* mRNA could accumulate in cytoplasmic foci for storage in response to heat stress. To assess this possibility, we analysed its localisation within the cell. The method chosen utilizes RNA loops (MS2SL) co-expressed with a coat protein (MCP), both derived from the bacteriophage MS2. Several MS2SL were inserted in the 3′ UTR of the endogenous *TPK1* mRNA by a gene-tagging procedure called m-TAG and MCP was expressed as a fusion protein with GFP (MS2-GFP). This system allows endogenous mRNAs in living yeast to be visualized (Haim-Vilmovsky and Gerst, 2009). The constructions were performed in a wild-type yeast strain (W3031A *DCP2-CFP eIF4E-RFP*). The heat-shock treatment was performed on exponential growing cells for 60 min at 37°C. Images obtained through confocal microscopy, showed that *TPK1* mRNA was distributed in discrete granules in the cells upon heat stress but not in non-treated cells (25°C) (Fig. 3A). Importantly, control cells containing the pMS2-CP-GFPx3 plasmid but lacking MS2SL on *TPK1*, exhibited diffuse GFP signal throughout their cytoplasm, showing that the MS2-GFP fusion *per se* does not produce any aggregation in the cells (Supplementary Figure 1). Some publications argue against the use of the MS2-GFP tool. Firstly, it has been reported that the introduction of MS2SL can affect mRNA processing, as their presence inhibits 5′ to 3′ degradation resulting in the accumulation of 3′ mRNA fragments containing MS2 binding sites (Garcia and Parker, 2015). Secondly, the transcripts containing MS2SL binding sites showed unusual localisation in glucose starvation conditions, with altered nuclear mRNA processing and stem–loop fragments enrichment in P-bodies (Heinrich et al., 2017). Accordingly, we performed Northern blot experiments to detect *TPK1* transcripts that contain MS2SL arrays using a specific probe directed to MS2SL sites. As 3′-end mRNA fragments, originated from mRNA degradation, were not detected in neither control nor heat shock treated cells and the mRNA detection was the same in both conditions (Supplementary Fig. 2), we can rule out that significant *TPK1* mRNA fragmentation occurs under our experimental conditions.

**Figure 3.**
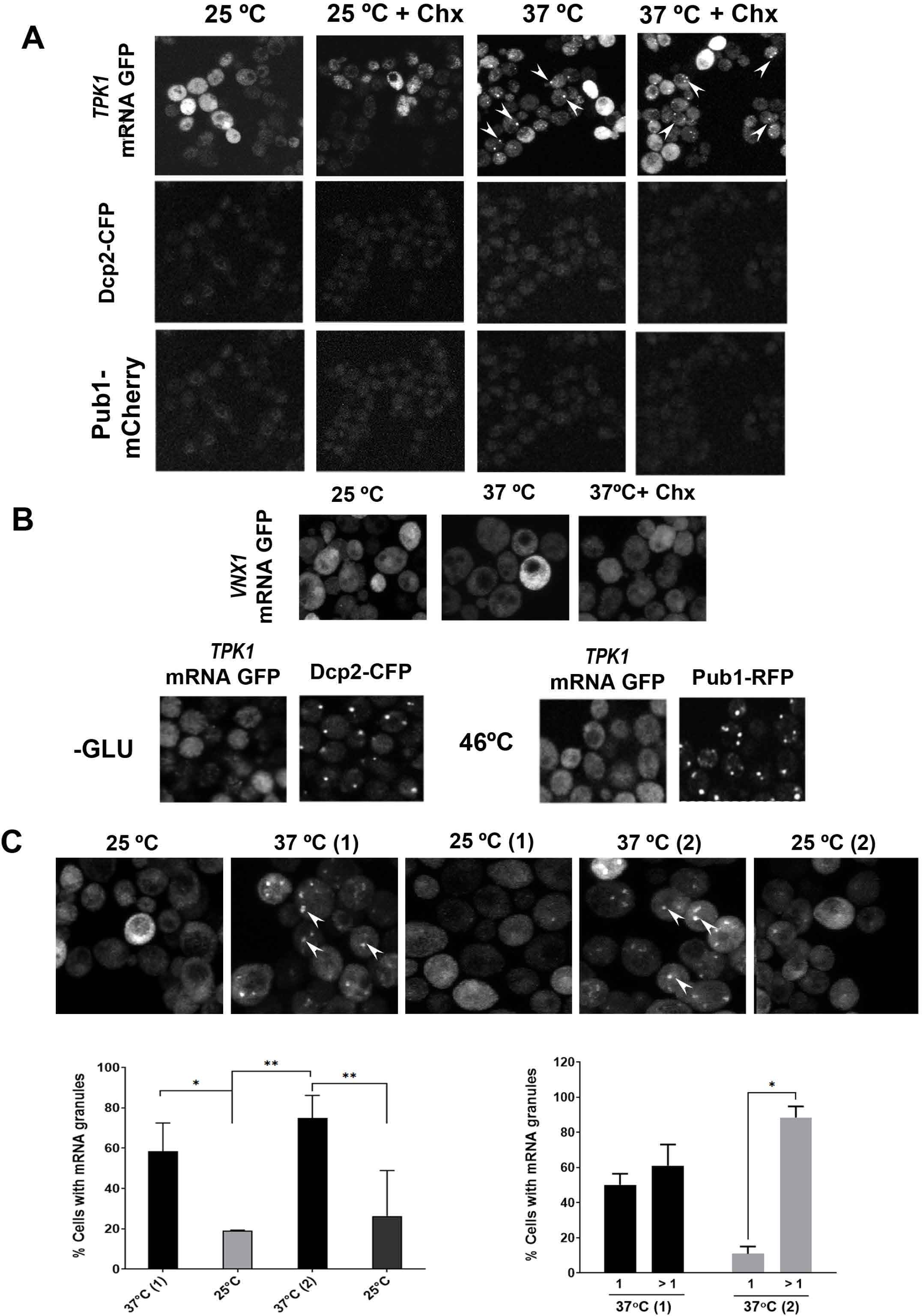
*TPK1* mRNA is assembled in foci upon heat shock. A) Cells co-expressing *TPK1* mRNA-MS2SL, pMS2-CP-GFP, Dcp2-CFP (P-bodies) and Pub1-mCherry (SGs) were incubated or not at 37°C for 60 minutes in the presence or not of cycloheximide. *TPK1* mRNA foci are indicated by arrowheads on panels. B) Cells co-expressing *VNX1* mRNA-MSLS2 and pMS2-CP-GFP were then incubated or not at 37°C for 60 minutes in the presence or not of cycloheximide (upper panels). Cells co-expressing *TPK1* mRNA-MSLS2 and pMS2-CP-GFP with Dcp2-CFP or Pub1-mCherry were deprived of glucose for 30 min or incubated at 46°C for 10 minutes, respectively (lower panels). C) *TPK1* mRNA foci kinetics through the heat shock adaptation. The left bar chart shows the percentage of cells with granules at the indicated temperatures and the right bar chart shows the percentage of cells with 1 granule or more than 1 granule per cell at the indicated temperatures. The values represent the mean ± s.e.m., obtained from n=4 biological replicates. Statistical analysis corresponds to two-way ANOVA, *p<0.05; **p<0.01.

To shed more light on the characteristics of the *TPK1* mRNAs granules, cycloheximide was used to block translational elongation. Cycloheximide cause polyribosome stalling and prevents the formation of cytoplasmic granules avoiding the movement of mRNA across the granules and leading to their disassembly (Anderson and Kedersha, 2009; Buchan et al., 2008; Grousl et al., 2009; Grousl et al., 2013; Kato et al., 2011; Teixeira et al., 2005). Cells were treated or not with cycloheximide before the heat shock and the presence of the *TPK1* mRNA granules was analysed (Fig. 3A). No changes in the number of granules per cell were observed after cycloheximide treatment suggesting that *TPK1* mRNA in the granules is prevented from entry to translation. Further characterization of the *TPK1* mRNA granules was assessed by analysing their colocalization with marker proteins of P-bodies or SG. Thus, the colocalization of heat shock responsive *TPK1* mRNA granules with CFP-Dcp2p (P-body component) or mCherry-Pub1p (SG marker) was assessed. In our experimental conditions, after 60 min at 37°C neither Dcp2 nor Pub1 foci were assembled, and this result suggests that the *TPK1* mRNA containing granules does not include these P-bodies or GS markers (Fig. 3A). As Dcp2p or Pub1p containing granules were not observed at 37°C 60 min, experiments to confirm the correct expression of these proteins were performed. Cells were subjected to known P-bodies and SG inducing conditions as glucose deprivation and severe heat shock at 46°C. Interestingly, even though the glucose deprived cells were efficient in P-body formation (Dcp2-GFP granules) and cells subjected to severe heat stress (46°C 10 min) formed SGs (Pub1-mCherry granules), *TPK1* mRNA granules were not detected under these conditions, and thus, colocalization with those markers could not be observed (Fig. 3B). As a control, the localization of another MS2-tagged mRNA was also assessed. It was previously described that *VNX1* mRNA localises into P-bodies at a late phase during glucose starvation (Simpson et al., 2014a). Thus, the localization of the *VNX1* was assessed in exponentially growing yeast subjected to 37°C or to 37°C plus cycloheximide. *VNX1* mRNAs were not detected in granules after these heat stress conditions but were distributed throughout the cytoplasm (Fig. 3B). This result indicates that the presence of the MS2-tag was not sufficient to drive mRNA granule assembly after the heat shock. Altogether, these results indicate that, upon mild heat stress, *TPK1* mRNAs associate to granules that do not contain Pub1p nor Dcp2p.

Next, the dynamics of *TPK1* mRNA granules during thermal adaptation were evaluated. The number of granules evoked during the first heat shock (58% of the cells contained granules), decreased after 40 min at 25°C (18% of the cells contained granules). After the second heat shock, the number of cells with *TPK1* mRNA granules increased (70% of the cells contained granules) and this number was again reduced after the second recovery period at 25°C (20% of the cells contained granules) (Fig. 3C, left bar charter). This *TPK1* mRNA localization pattern correlates with the variations in protein production and mRNA abundance detected upon heat shocks and recoveries (Fig. 2A). Western blot analysis in both BY4741 and TPK1-MS2SL strains showed similar Tpk1 expression patterns (data not shown).

The number of granules formed per cell varied between the first and the second heat shock. The second 37°C incubation evoked more *TPK1* mRNA foci per cell than the first one (Fig. 3C, right bar charter).

All these results suggest that the *TPK1* mRNAs upregulated upon thermal stress could be translationally silenced and stored in mRNP granules. Furthermore, the fact that Tpk1 levels during the second incubation at 25°C are higher than those during the first one, possibly represents a stress adaptation mechanism.

### *TPK1* mRNA granules in heat-stressed cells can be associated with repressed translation

There are a few examples of cycloheximide resistant foci that contain mRNA. In response to α-pheromone, *MFA2* mRNA is assembled in two types of mRNA granules. The smaller ones contain untranslatable *MFA2* mRNA and are involved in mRNA transport to the tip of the shmoo contributing to the localised translation of *MFA2* (Aronov et al., 2015). TT foci, present in the cytosol of THO/TREX-2 yeast mutant cells, are also cycloheximide resistant and contain aberrant exported mRNPs (Eshleman et al., 2016). For both examples, the authors concluded that the mRNAs in these foci are translationally incompetent. Taking into account these antecedents and the discrepancy between the levels of both steady state *TPK1* mRNA and Tpk1 protein during heat stress, we understand that the unsensitivity of *TPK1* mRNA foci to cycloheximide treatment could be a consequence of *TPK1* mRNA being incompetent for translation.

Thus, we performed polysome profiling followed by RT-qPCR to determine the levels of the *TPK1* mRNA associated with polysome (P) and sub-polysome fractions (M and F). Figure 4A shows the polysome analysis performed in exponentially grown wild type cells without MS2-tagging (WT-A) incubated at 25°C and 37°C. Around 40% of the *TPK1* mRNA in untreated cells was present in the polysome fraction. After heat stress, this proportion declined up to 20% with a concomitant increase of *TPK1* mRNA in sub-polysome fractions (Fig. 4A). The quality of the approach was controlled by analysing the presence of specific major and minor ribosomal subunit proteins by western blot (Fig. 4B). The correct distribution of these proteins in the fractions confirms that the polysome profile is suitable. The distribution of *HSP30* and *ENO2* mRNAs, whose translation is upregulated upon heat shock, in polysome or sub-polysome fractions was also evaluated. Hsp30 is a negative regulator of the H (+)-ATPase Pma1p and a stress-responsive protein induced by heat shock (Barraza et al., 2017). The *HSP30* mRNA amount in the polysome fraction increased upon heat stress (Fig. 4C, right panel). *ENO2* encodes the enolase II protein and its translational activity decreased upon heat stress (Barraza et al., 2017)(Fig. 4C, bottom panel). The distribution of these mRNAs in P and SP fractions agrees with their increase or decrease protein levels upon heat shock. Finally, the distribution of *BCY1* mRNA to each polysome fraction was also analysed and it did not change with stress (Fig. 4C, left panel). This distribution agrees with the constant levels of the Bcy1 protein during heat shock. All these results indicate that the translation of *TPK1* mRNAs is less active upon heat shock.

**Figure 4.**
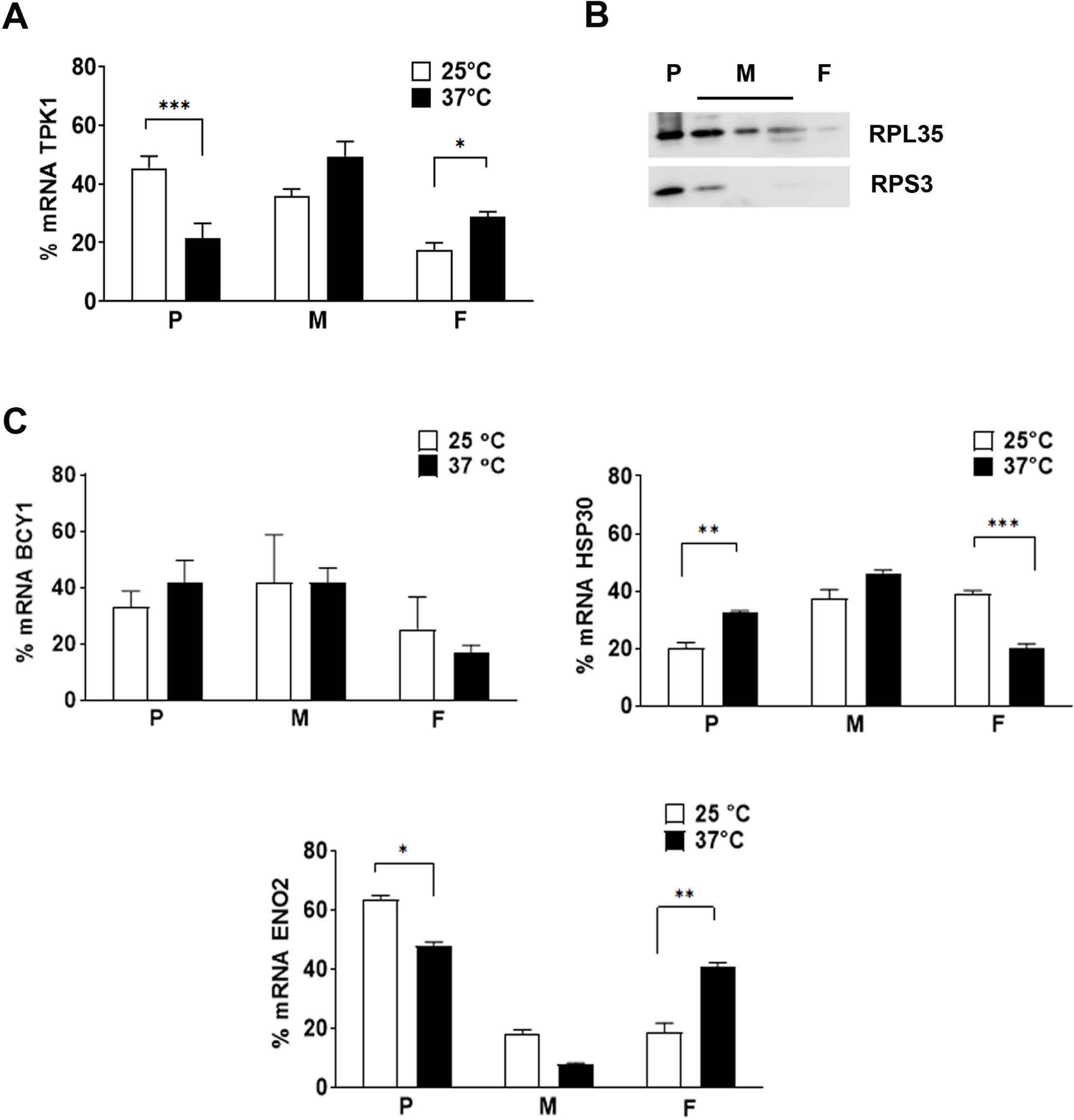
*TPK1* mRNA is less translated upon heat shock. A) Polysome profile and RT-qPCR analysis of RNA prepared from individual fractions of polysome gradients from cells treated or not for 60 min at 37°C. The bar chart represents the percentage of *TPK1* mRNA abundance in the sub-polysome and polysome fractions. The values represent the mean ± s.e.m. from n=4 biological replicates. Statistical analysis corresponds to two-way ANOVA, *p<0.05; **p<0.01; ***p<0.001. B) Fractions were also analyzed by SDS–PAGE and immunoblotting using antibodies raised against the proteins specified adjacent to each panel. P (polysome fraction), M (Monosome fraction, 40S, 60S and 80S), F (free RNA). C) Polysome distribution of *BCY1* mRNA, *HSP30* mRNA and *ENO2* mRNA as described in A.

### *TPK1* expression is regulated through the CWI pathway

Several signalling pathways communicate at different levels to integrate different types of stress. This crosstalk enables fine-tuned regulation of stress responses. The cAMP-PKA pathway may be connected to other signalling stress sensing pathways to ensure yeast cells survival. The CWI pathway regulates the cell wall biosynthesis upon several stresses including temperature shifts (Donlin et al., 2014; García et al., 2017; Heinisch, 2020; Heinisch and Rodicio, 2009; Petkova et al., 2010)Fuchs and Mylonakis, 2009; Heinisch, 2020; Levin, 2011; Sanz et al., 2017). We wondered whether crosstalk between the CWI and cAMP-PKA signalling pathways is involved in controlling the *TPK1* expression in response to heat stress. The activation of the CWI pathway by heat shock and the participation of the different Wsc receptors in this response have been demonstrated (Levin, 2011; Verna et al., 1997; Zu et al., 2001). The activity of *TPK1* promoter using a promoter-lacZ based reporter assay and the levels of *TPK1* mRNA were assessed in yeast strains with deletions in the cell surface sensors *wsc1Δ, wsc2Δ, wsc3Δ* (Fig. 5A and B), *mid2Δ* and *mtl1Δ* (Supplementary Fig. 3). The results show that the transcriptional induction of *TPK1* upon heat shock depended only on the Wsc3 sensor (Figure 5A and 5B; Supplementary Figure 3 A and B). The Tpk1 protein levels in *wsc3Δ* cells upon thermal stress were also analysed by western blot. The levels of Tpk1, at t=0 and after the heat shock, were higher in the *wsc3Δ* than in the WT strain (Fig. 5C). Heat shock induces the activation of Slt2/Mpk1 and therefore an increase in its phosphorylation (González-Rubio et al., 2021; Jiménez-Gutiérrez et al., 2020; Martín et al., 2000; Zarzov et al., 1996). We assessed the activation of CWI pathway and the involvement of Wsc3 in our stress conditions, evaluating Stl2 phosphorylation by western blot. Figure 6A shows that the Slt2 phosphorylation in the WT strain, but not in the *wsc3Δ,* significantly increased in response to the heat shock.

**Figure 5.**
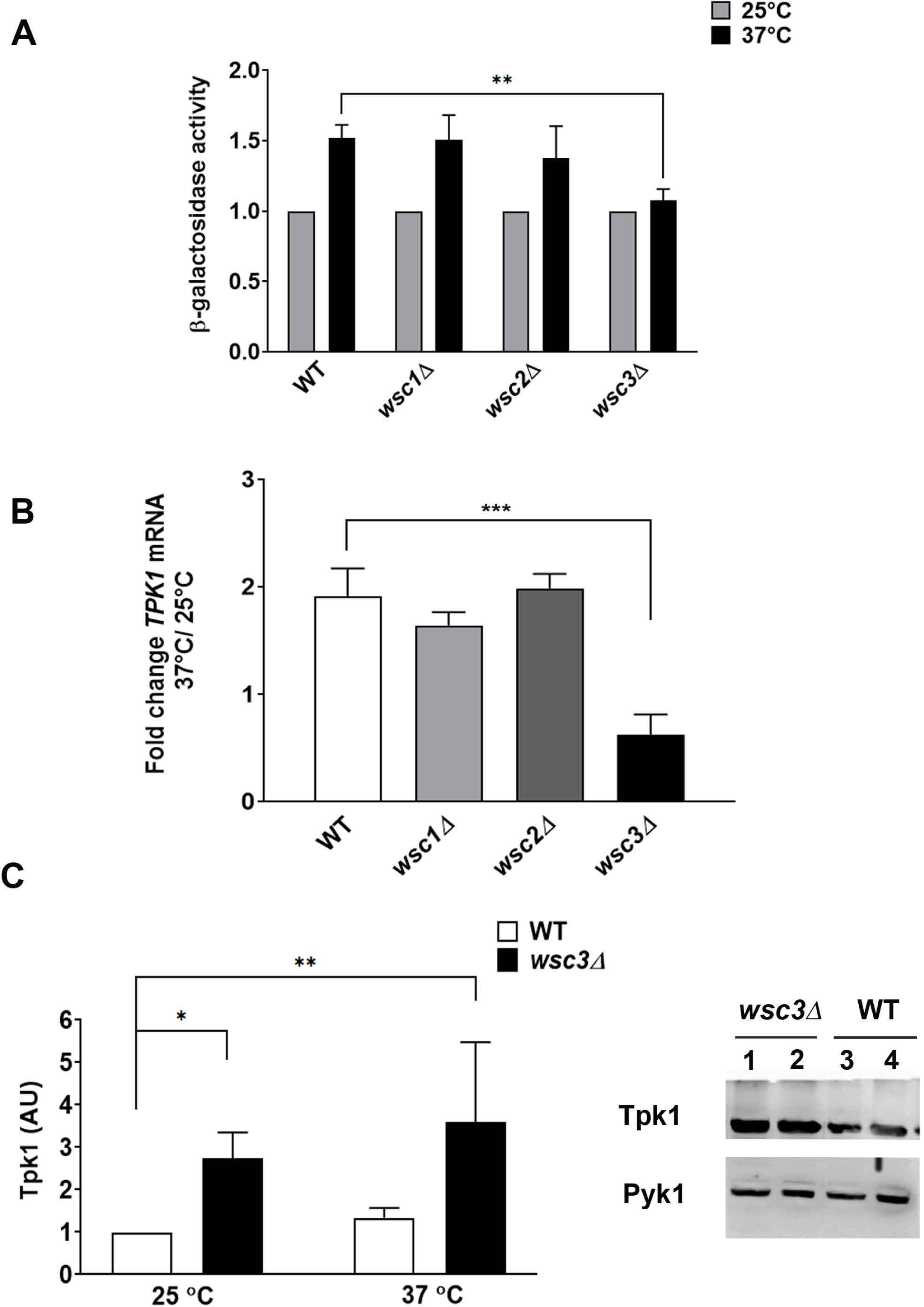
*TPK1* mRNA upregulation is dependent of wsc3 sensor. A) β-Galactosidase activity was determined in WT (BY4741), *wsc1Δ, wsc2Δ* and *wsc3Δ* strains carrying *TPK1-*lacZ fusion gene incubated or not at 37°C for 60 min. The results are expressed in Miller Units and as the mean ± s.e.m. of biological replicates samples (n= 4). B) *TPK1* mRNA levels were determined by RT-qPCR from WT, *wsc1Δ, wsc2Δ* and *wsc3Δ* cells and normalized to *TUB1* mRNA and expressed as 37°C/25°C fold change. The values represent the mean ± s.e.m., obtained from n=3 biological replicates. C) Tpk1 protein levels were analyzed by western blot in protein extracts from WT and *wsc3Δ* cells incubated or not at 37°C for 60 min using anti-Tpk1 antibody. The values represent the mean ± s.e.m., obtained from n=3 biological replicates. Statistical analysis corresponds to two-way ANOVA, *p<0.05; **p<0.01; ***p<0.001

**Figure 6.**
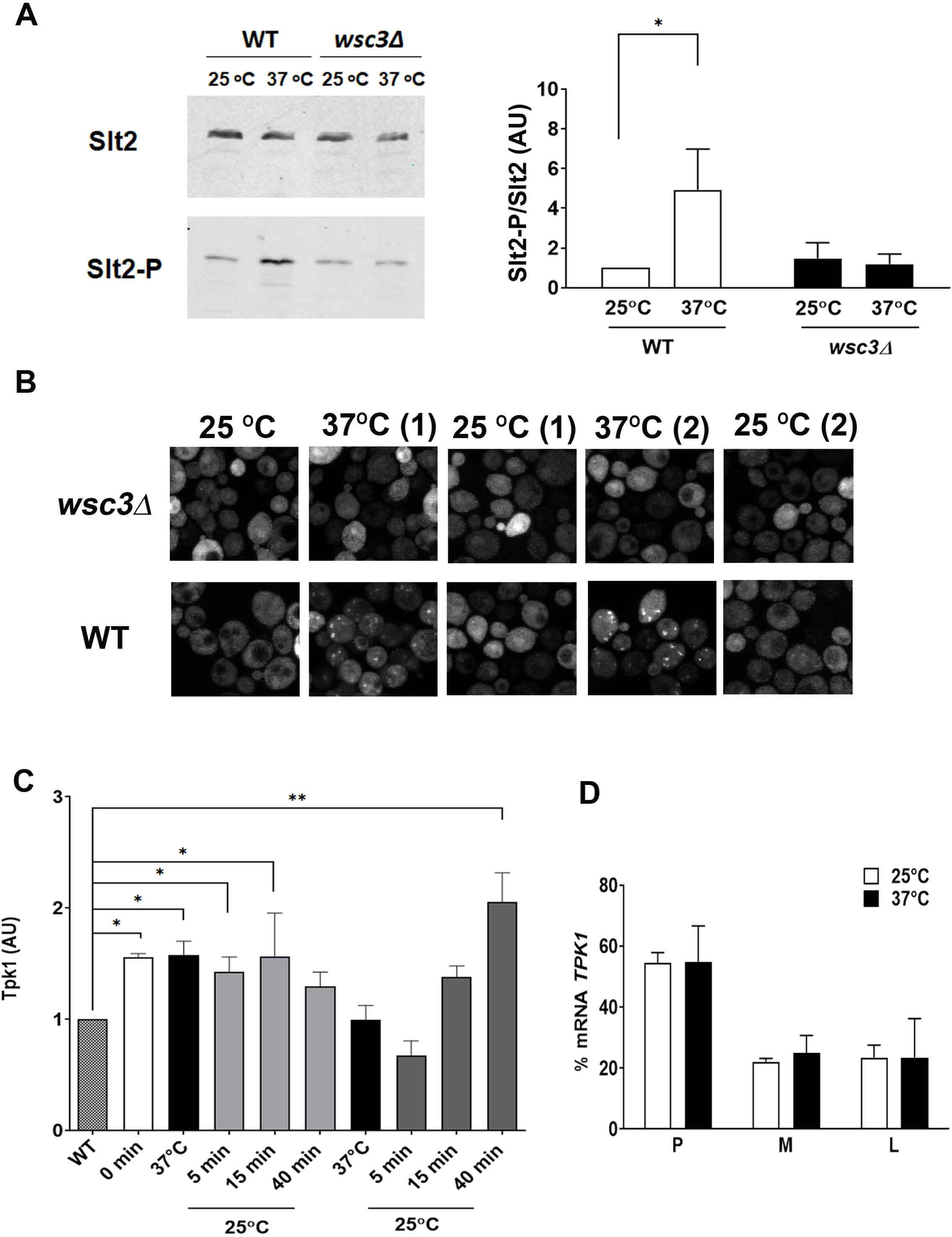
*TPK1* mRNA foci evoked upon heat shock are dependent of wsc3 sensor. A) Extracts from the WT and *wsc3Δ* cells incubated or not at 37°C for 60 min were analysed by Western blot with Slt2-p antibody (phosphorylated) or Slt2 (total protein), quantified using Image J and expressed as AU (arbitrary units, right panel). B) WT (W303-1A) or *wsc3Δ* cells co-expressing mRNA-MSLS2 and pMS2-CP-GFP were incubated or not at 37°C for 60 minutes. *TPK1* mRNA foci are indicated by arrowhead on panels. C) *wsc3Δ* cells were treated following the scheme presented in Fig 2A. Samples were analysed by Western blot using anti-Tpk1. The bar chart represents the quantification of Tpk1 levels expressed in AU (arbitrary unit, left panel). The values represent the mean ± s.e.m., obtained from n=4 biological replicates. D) Polysome profile and RT-qPCR analysis of RNA prepared from individual fractions of polysome gradients from wsc3Δ cells. The values represent the mean ± s.e.m., obtained from n=4 biological replicates. Statistical analysis corresponds to two-way ANOVA, *p<0.05; **p<0.01; ***p<0.001.

We showed above that, in heat stressed wild type cells, *TPK1* mRNA is assembled in untranslated *TPK1* mRNA granules. Since a role of Wsc3 in the regulation of *TPK1* expression was demonstrated, the localization of *TPK1* mRNA using the MS2-tagged endogenous mRNA in a *wsc3Δ* strain was also analysed. Cells were subjected to the scheme of temperature shifts described before (Figure 2A). In a *wsc3Δ* strain, *TPK1* mRNA localisation was altered compared to WT strain upon heat stress. Diffuse distribution of *TPK1* mRNA was observed throughout the cytoplasm in the mutant strain (Fig. 6B). Thus, *TPK1* mRNA foci, observed in heat stressed wild-type cells were dependent on the Wsc3 sensor. Changes in Tpk1 protein levels throughout the experiment were also analysed (Fig. 6C). Initial Tpk1 levels (time 0) were higher in the *wsc3Δ* than in the WT strain and these levels did not increase after thermal stress (Fig. 5C and 6C). Only during the second 25°C incubation, the protein levels detected in *wsc3Δ* cells were significantly higher than the observed before any stress. It is noteworthy that the increase was relatively small compared to that observed in wild-type cells (Fig. 2B and 6C). The results indicate that the Wsc3 sensor is involved in the regulation of *TPK1* expression during heat shock and stress adaptation. The translation of *TPK1* mRNA in *wsc3Δ* cells was assessed by analyzing *TPK1* mRNA levels associated with polysome and sub-polysome fractions before and upon heat stress. The amount of *TPK1* mRNA in polysome fractions was higher than that observed in the same fraction in wild-type cells (55% versu*s* 40%, Fig. 6D and Fig. 4A), but the thermal stress did not alter the polysome profiles in the mutant (Fig. 6D). These results suggest that the *TPK1* mRNA, though less abundant, may be more actively translated in *wsc3Δ* cells.

Some studies indicate that the signal that activates Slt2 in the CWI pathway is not transduced in a simple top–down cascade form since the CWI sensors showed to be dispensable for Slt2 phosphorylation in response to some stresses (Harrison et al., 2001; Krause and Gray, 2002), such as heat shock (Harrison et al., 2004). It was demonstrated that heat-shock triggers this signalling pathway regulating either Mkk1/2p or Slt2/Mpk1p activity directly without the participation of Rho1p GTP (Harrison et al., 2004; Kuravi et al., 2011). We analysed the involvement of different CWI pathway components as Rom2, Pkc1, the downstream MAP kinase cascade, Bck1, Mkk1 and Slt2 in *TPK1* expression using the *TPK1*-lacZ reporter plasmid system and the quantification of the mRNA levels (Supplementary Fig. 3B and C). The results showed that Mkk1 and its downstream effector, the transcription factor Swi4, but not Swi6, regulated *TPK1* transcription. Intriguingly, although the heat shock-induced phosphorylation of Slt2 was impaired in the *wsc3Δ* mutant, *TPK1* expression was not affected by the lack of this kinase (Supplementary Fig. 3B and C). There are antecedents of genes whose expression is dependent on Swi4 but independent on both Swi6 and Slt2. Moreover, a little overlap was described between *swi4Δ* and *stl2Δ* gene profiles after heat shock (Baetz et al., 2001). Thus, together with these antecedents, our results support a unique and specific role of the CWI pathway in *TPK1* expression.

### Wsc3 modulates PKA activity *in vivo*

Having established a connection between the sensor Wsc3 and *TPK1* expression, the *in vivo* PKA activity in *wsc3Δ* strains was analysed. We assessed some physiological readouts of the cAMP-PKA pathway and PKA activity, such as heat resistance, glycogen accumulation and intracellular distribution of Msn2 (Cameron et al., 1988; Görner et al., 1998; Shin et al., 1987; Smith et al., 1998; Thevelein and Winde, 1999). The mutant *wsc3Δ* was less resistant to a 52°C-heat shock than the wild-type strain indicating a higher PKA activity in the mutant strain (Supplementary Fig. 4A). This result agreed with a higher level of Tpk1, detected by western blot, in the mutant strain than in wild-type cells when they were unstressed (Fig. 6C). In unstressed *wsc3Δ* cells this higher PKA activity was also confirmed by the lower glycogen accumulation observed (Supplementary Fig. 4B, left panel). Glycogen accumulation in cells from both recovery periods at 25°C, after each heat shock, was assessed. *wsc3Δ* strain showed lower glycogen levels than wild-type cells (Supplementary Fig. 4, right panel). This result suggests a lower PKA activity in the mutant than in the wild-type strain during the recovery periods at 25°C.

Since high PKA activity leads to nuclear Msn2 redistribution to the cytoplasm, the nuclear/cytoplasmic localisation of Msn2 in WT and *wsc3*Δ cells expressing Msn2-GFP was assessed under control and heat shock conditions. The results indicate that the Msn2 localization kinetics were similar in both strains (Supplementary Fig. 4C). The translocation of Msn2 to the nucleus occurred within the first 5 min of the first heat shock and it was transient. However, the percentage of *wsc3*Δ cells with nuclear localization of Msn2 through the temperature shifts was lower than that detected in the wild type except for the unstressed cells (t=0) (Supplementary Fig. 4C). These results suggest that, during the recovery periods, PKA activity is higher in wild-type than mutant cells. Collectively, all these results demonstrate that the Wsc3 sensor is involved in the modulation of PKA activity *in vivo* when yeast cells are exposed to heat shock.

Finally, we performed experiments to assess whether the increment in Tpk1 protein upon heat shock results in a higher proportion of the Tpk1 catalytic subunit in the PKA holoenzyme. The rationale behind this new approach is based on the idea that the differential expression of Tpk1 isoform may contribute to the cAMP-PKA signalling specificity. Yeast cells were treated with 2% formaldehyde to cross-link complexes. Lysates were incubated at 65°C and 100°C and analyzed by western blot using anti-Tpk1 and anti-Bcy1 antibodies (Kast and Klockenbusch, 2010). As formaldehyde cross-linking is reversed by boiling, cross-linked complexes would be hardly detected in the samples heated at 100°C. After heating at 65°C, cross-linked complexes are much less disassembled and would be detected. Figure 7A shows that in the sample treated with 2% formaldehyde and 65°C complexes containing Bcy1 were detected. Figure 7B shows the Tpk1-Bcy1 complexes detected in extracts obtained from yeast grown at 25°C (line 1) or from the first (line 2) and second recovery periods (line 3). The amount of Tpk1 relative to Bcy1 in the holoenzyme was higher in the second 25°C recuperation period than in the first one or before any thermal stress (Fig. 7B). However, the amount of Bcy1 does not change in the samples at the recuperation periods (Figs. 2 and 7B). The amount of Tpk1 in purified holoenzyme was also assessed using tandem affinity purification (TAP) assay and a strain expressing the fusion protein Bcy1-TAP. The presence of Tpk1 and Bcy1 subunits in the purified holoenzyme complex was evaluated by immunoblotting (Fig.7 C). Even though the differences between Tpk1 amounts in the control and after the second recuperation period were not statistically significant, the results show the same tendency observed in the crosslinking assays.

**Figure 7.**
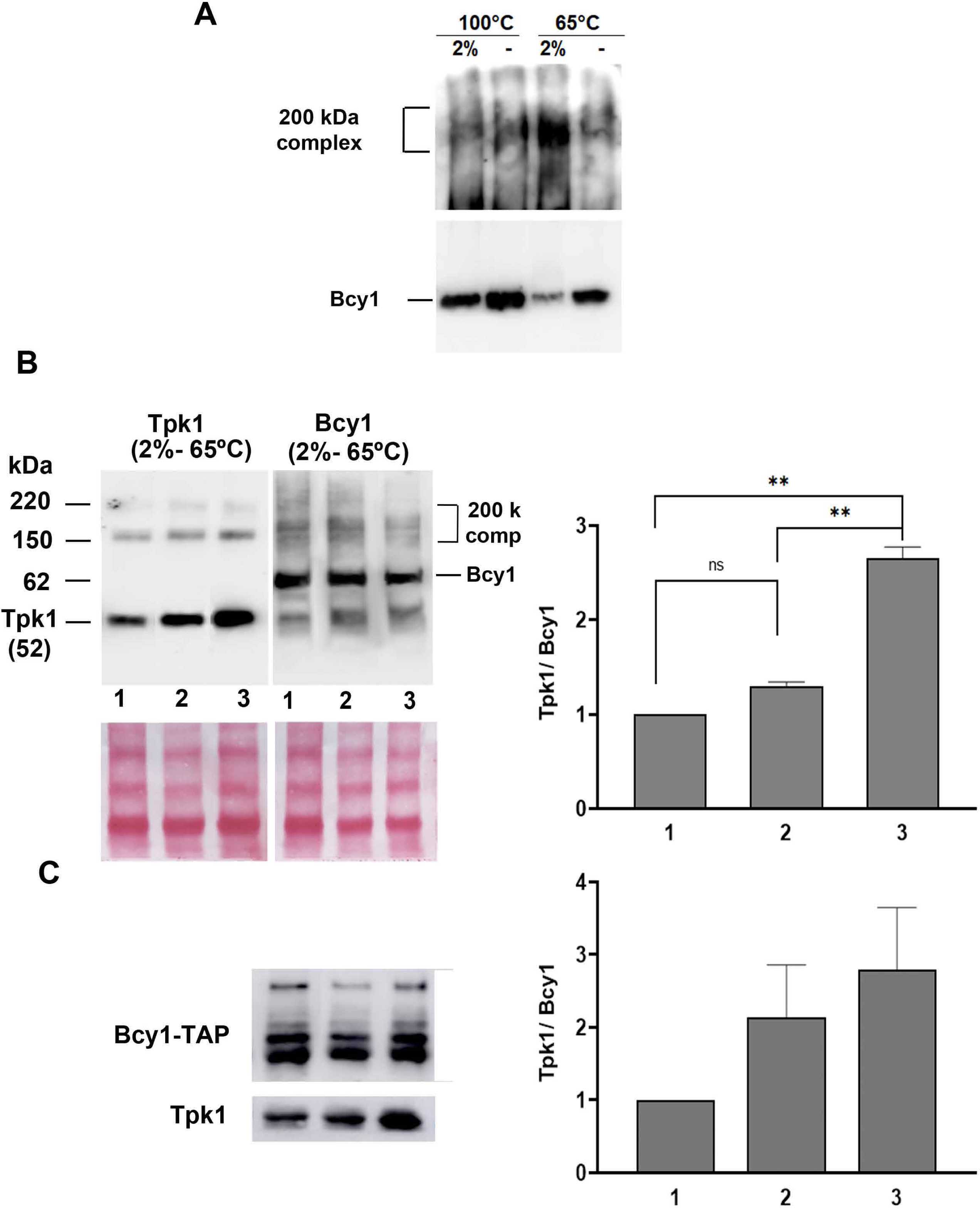
PKA holoenzyme is composed by a higher proportion of Tpk1 isoform upon heat shock. A) Cells were crosslinked or not with formaldehyde 2%. Cell extracts were prepared, heated at 100°C or 65°C and analysed by Western blot using anti-Bcy1. Bcy1 crosslinked complexes were overexposed (upper panel). B) Cells were treated following the scheme presented in Fig 2A. The samples taken out at t=0 and during the recovery periods were crosslinked with formaldehyde 2%. Cell extracts heated at 65°C were analysed by Western blot using anti-Bcy1 or anti Tpk1. Line 1, t=0; line 2, 25°C (1); line 3, 25°C (2). Protein molecular weight markers are indicated. Bottom panel: Ponceau staining was used as loading control. Right panel: quantification of Tpk1/Bcy1 in the holoenzyme relative at t=0, 25°C (1) and 25°C (2). C) The amounts of Tpk1 and Bcy1 from a purification of Tpk1-Bcy1-Tap were analyzed by SDS-PAGE following immunoblot with anti-Tpk1 and anti-Bcy1 (left panel). The graph shows the quantification of Tpk1/Bcy1 in the purified holoenzymes (right panel).

These results suggest that the induced expression of Tpk1 during adaptation to thermal stress results in PKA holoenzymes that contain a greater proportion of the Tpk1 isoform which would assure the phosphorylation of its specific substrates.

## DISCUSSION

Several levels of control acting simultaneously and in coordination allows the specificity of the cAMP-PKA signal transduction. Our previous results have demonstrated that the mechanisms involved in *TPK*s and *BCY1* expression in response to stress or growth conditions are subunit-specific (Galello et al., 2017; Pautasso and Rossi, 2014; Reca et al., 2020).

Yeast cells respond to a variety of environmental stresses, such as heat shock, by reprogramming the expression of specific sets of genes for any particular stress (Berry and Gasch, 2008; Gasch et al., 2000; Guan et al., 2012; Molina-Navarro et al., 2008; Morano et al., 2012; Rienzo et al., 2015). Studies on stress responses demonstrate that mRNA levels result from the balance between the transcription rate and the decay rate of each particular mRNA (Pérez-Ortín et al., 2019). Through this study, we were successful in further understanding the mechanisms involved in the regulation of *TPK1* expression during heat shock. The analysis of mRNA half-lives showed that t1/2 of *BCY1* mRNA is higher than that of *TPK1* mRNA (19.5 min versus 4.6 min) at control conditions., However, both *BCY1* and *TPK1* mRNA t1/2 are increased upon heat shock. The assessment by western blot of protein levels during the first and second heat shocks showed that there were no changes in Tpk1 levels after the incubations at 37°C, although an increase in protein levels was observed during the incubations at 25°C (first and second recovery periods). The levels of Bcy1 protein showed no significant changes even though the mRNA half-life was greater at 37°C than at 25°C. It has been previously reported that the half-life values of Tpk1 and Bcy1 proteins remain unchanged after a heat shock at 37°C (Budhwar et al., 2010; Christiano et al., 2014, Haesendonckx et al., 2012). Thus, the lack of changes in Bcy1 and the variations in Tpk1 levels at the evaluated temperatures do not seem to result from differences between the protein half-life values.

The results obtained in the evaluation of mRNA localization allow us to conclude that the increased *TPK1* mRNA is assembled in foci upon heat shock. Recently, some reports have pointed out that the study of mRNA localization using MS2-tethering techniques should be approached with caution when interpreting data, since the MS2 stem loops may stabilize mRNA fragments and alter the RNA processing (Garcia and Parker, 2015; Garcia and Parker, 2016; Haimovich et al., 2016; Heinrich et al., 2017). However, it has also been demonstrated that this phenomenon is limited to a subset of endogenous MS2-tagged transcripts and associated with plasmid-based expression systems (Haimovich et al., 2016). We have performed Northern blot assays to analyse the possible accumulation of 3’end fragments from *TPK1* mRNA tagged with the MS2SL–MCP system. The results showed that *TPK1* mRNA, from both heat shock and control conditions, did not lead to the formation of 3′ decay fragments containing MS2 arrays, and in both conditions the integrity of the mRNA are equivalent. We conclude that the addition of MS2SL to the *TPK1* mRNA does not affect the processing nor stability of the mRNA as it has been previously observed for other mRNAs (Simpson et al., 2014). In addition, the increased *TPK1* mRNA half-life upon heat shock was assessed using endogenous mRNA without MS2SL-tagging and oligo probes complementary to the *TPK1* CDS, in the middle of the coding sequence. The increased abundance of the *TPK1* mRNA upon heat shock, assessed by RT-qPCR, was performed analysing endogenous *TPK1* mRNA without MS2SL-tagging. Finally, the measurement of Tpk1 protein levels throughout the scheme of consecutive heat shocks was assessed to the two strains, WT-A and *TPK1*-MS2SL, and the observed expression pattern was the same in both (data not shown). Thus, we consider that our results of *TPK1* mRNA foci kinetics are valid and not artifacts produced by the tagged mRNA-MS2SL technique.

The *TPK1* mRNA foci evoked upon heat shock differ from other cytoplasmatic RNA granules. The composition and assembly of RNP granules as P-bodies and SGs are complex and appear to vary in a stress-dependent manner (Guzikowski et al., 2019; Thomas et al., 2011). Most studies describe that SGs are assembled at severe but not at mild heat stress conditions (Grousl et al., 2009; Grousl et al., 2013; Kato et al., 2011). Dcp2-CFP positive foci were detected after incubation at 37°C for 10 min or 30 min (Barraza et al., 2017; Brengues et al., 2005; Grousl et al., 2009; Grousl et al., 2013). Under our conditions (37°C 60 min), neither P-bodies nor SGs were detected. *TPK1* mRNA foci were assembled upon the mild heat shock conditions assessed in this work, but they were not evoked under severe heat shock conditions. Taking into account these results we conclude that *TPK1* mRNA foci constitute another class of RNA granule that differs from P-bodies and SGs. Granules that differ from P-bodies and SGs may be assembled through a distinct biological process or different stresses and these structures may be composed of particular proteins. In stationary-phase yeast cells, proteins associated with the actin cytoskeleton and the proteasome are found in actin bodies and proteasome storage granules, respectively (Laporte et al., 2008; Sagot et al., 2006). RNP granules also appear when yeast cells stop dividing and become quiescent (Narayanaswamy et al., 2009; Noree et al., 2010).

The SG and P-body assembly in yeast cells is avoided by cycloheximide which causes the stalling of polyribosomes on mRNA (Brengues et al., 2005; Buchan et al., 2008). However, mRNA *TPK1* granules were resistant to cycloheximide treatment. We demonstrated that, while these foci are present, *TPK1* mRNA associates more to subpolisome (M plus F) than to polysome fractions. We conclude that *TPK1* mRNA less translated upon thermal stress, which agrees with the lack of Tpk1 protein increase upon heat stress. There are examples of cycloheximide resistant foci, such as the smallest *MFA2* mRNA granules which are not translated (Aronov et al., 2015) and the TT foci related to the THO/TREX-2 complex (Eshleman et al., 2016). Low correlations between steady state mRNA and protein levels have been shown, which is a consequence of substancial post-transcriptional regulation (Csárdi et al., 2015; Wu et al., 2008). We have demonstrated that Tpk1 levels upon heat stress are modulated by a post-transcriptional mechanism that involves the assembling of translationally silent *TPK1* mRNA granules. During recurrent thermal stress the changes in *TPK1* mRNA assembling and Tpk1 levels shows that yeast cells regulate the *TPK1* expression. The final result is a high level of Tpk1 protein at 25 °C when yeast cells are not exposed to the stress.

Our results demonstrate that CWI pathway is involved in Tpk1 expression. The involvement of different CWI sensors in response to environmental changes has been established. None of the single sensor mutants *wsc1Δ*, *wsc2Δ*, or *wsc3Δ* are thermosensitive, while combinations of these mutants show a thermosensitive phenotype (Verna et al., 1997). Later on, it was concluded that Wsc proteins produce an additive effect in response to specific stresses as heat shock (Zu et al., 2001). It has been reported that some stress events can regulate this pathway at different steps downstream Rho1-GTP. Heat shock activates the CWI pathway at the level of Mkk1/2p and/or Mpk1p (Harrison et al., 2004). This study demonstrates that the Wsc3 sensor and Mkk1, but not Mpk1, are necessary to activate the *TPK1* promoter upon heat shock. The polysome profiles of wild type and *wsc3Δ* cells were similar (data not shown), which indicates that the mutation does not affect the general translation process. However, Tpk1 translation upon thermal stress in *wsc3Δ* mutant was different that in the wild type. The transcription factor Swi4 also seems to be necessary for the regulation of the *TPK1* promoter. However, we are not able to distinguish the role of Swi6 since different results were obtained when *TPK1* promoter activity and mRNA levels were assessed. It has been described that there is little overlap between *swi4Δ* and *slt2Δ* gene profiles upon heat shock, and that there are genes whose expression is dependent on Swi4 but independent on both Swi6 and Slt2 (Baetz et al., 2001). Finally, although phosphorylation of Slt2 was diminished in the *wsc3Δ* strain upon heat shock, this kinase does not appear to play a role in the positive regulation of TPK1 by heat stress.

All these results indicate the involvement of the CWI pathway in *TPK1* expression upon heat stress. CWI pathway may be activated from Mkk1 of the MAPK cascade, resulting in an appropriate PKA output. Wsc3 would be the sensor of this lateral input in a way that seems independent of the activation of upstream elements of the CWI route. In fact, Rom2 does not seem to be involved either. This highlights the flexibility of the CWI signalling pathway and a crosstalk between CWI and PKA vias.

While trying to understand the implication of heat stress on the conformation of PKA holoenzyme, the levels of Tpk1 relative to Bcy1 were studied. Tpk1 proportion increased during both recovery periods compared to the amount observed before the stress.

We have previously proposed the existence of an apparent paradox during stress: *TPK1* PKA subunit transcription is stimulated during stress, despite higher PKA activity leading to lower stress resistance. The increment in Tpk1-dependent PKA activity during the adaptation of cells to heat stress could contribute to the overall cellular fitness when more favorable environmental conditions are restored. Our findings support the idea that the specificity in the cAMP-PKA signalling also results from differences in the PKA holoenzyme composition.

## MATERIALS AND METHODS

### Yeast strains, plasmids, culture conditions

S. cerevisiae strains and plasmids used in this study are summarized in Supplementary Table 1 and Table II, respectively. The plasmid used to measure promoter activity is derived from YEp357 plasmid (Myers et al., 1986). The TPK1-lacZ fusion gene contains the 5’ regulatory region and nucleotides of the coding region of TPK1 gene (positions -800 to +10 to the AUG). Strains were grown to mid log phase in synthetic media (SD) containing 0.67% yeast nitrogen base without amino acids, 2% glucose plus the necessary additions to fulfill auxotrophic requirements (SC). MS2 binding sites (MS2SL) were inserted into the 3’UTR of TPK1 gene and were verified using PCR and RT-PCR (Haim et al., 2007). Cells were grown at 25°C until exponential phase (OD600: 0.8-1). For heat shock treatment exponential cells were exposed to 37°C for 1 hour or the indicated times.

### RT-qPCR

Total RNA was prepared from different yeast strains, grown up to OD600 0.8-1 at 25°C and subjected or not (control) to heat shock (60 min at 37°C), using standard procedures. To determine the relative levels of specific TPK1 mRNAs, a quantitative RT-PCR experiment was carried out. Aliquots (∼10 μg) of RNA were reverse-transcribed into single-stranded complementary cDNA using an oligo-dT primer and M-MLV Reverse Transcriptase (Promega). The single-stranded cDNA products were amplified by PCR using gene-specific sense and antisense primers (mRNA TPK1: Fw: 5′ CCGAAGCAGCCACATGTCAC 3′, Rv: 5′ GTACTAACGACCTCGGGTGC 3′; mRNA BCY1: Fw: 5′ CGAACAGGACACTCACCAGC 3′, Rv: 5′ GGTATCCAGTGCATCG GCAAG 3′; mRNA TUB1: Fw: 5′ CAAGGGTTCTTGTTTACCCATTC 3′, Rv: 5′ GGATAAGACTGGAGAATATGAAAC 3′; mRNA ENO2: FW 5’ GAGAATCGAAGAAGAATTGGGTG3’, Rv: 5’CTATTCGTTAATATAAAGTGTTCTAAACTATGATG 3’; mRNA HSP30: Fw 5’ GTCTAAGTGATGGTGGTAAC 3’, Rv 5’ CTAAGCAGTATCTTCGACAG 3’). The PCR products were visualized using SYBR Green. The real-time qPCR reactions were performed on StepOne system (Applied Biosystems). The reactions were performed in duplicates and the results were standardized with the endogenous control of TUB1 (α-Tubulin gene). Quantitative data were obtained from three independent experiments and averaged.

### Immunoblotting

Crude extracts from cell grown up to OD 600 0.8-1 were separated by SDS-PAGE and transferred to GE Health Care or Osmonics nitrocellulose membranes. The blots were probed with anti-Tpk1 (Santa Cruz Biotechnology) and anti-Pyk1 (Polyclonal anti-pyruvate kinase (rabbit muscle) from Rockland) antibodies. Horseradish peroxidase-conjugated with anti-rabbit HRP and anti-mouse HRP (Sigma) were the secondary antibodies used. The blots were developed with prepared Chemiluminescence Luminol reagent (Sigma). The images were analysed by digital imaging using Amersham Imager 600. The quantification was performed using Image J and expressed as AU (arbitrary units).

### β-Galactosidase assays

Cells were grown on SC medium up to an OD600 of 0.8-1. Aliquots (10 ml) of each culture were collected by centrifugation and resuspended in 1 ml buffer Z (60 mM Na2HPO4, 40 mM NaH2PO4, 10 mM KCl, 1 mM MgSO4). β-Galactosidase activity measured according to Miller was expressed as Miller Units (Miller JH, 1972).

### Northern-blot and mRNA half-life analysis

Yeast cell cultures were grown in SC medium up to OD600 0.8–1 at 25°C and subjected or not (control) to heat shock (37°C) for the indicated times. 1,10-phenanthroline (Sigma, P-9375) was added to the cells to a final concentration of 100 µg.ml-1. For northern-blot analysis, aliquots were collected at the indicated times and the total RNA was extracted by the hot phenol method. Total RNA was separated in 1.5% agarose plus 6,7 % formaldehyde, transferred onto a nylon membrane and hybridized with 32P-labelled probes. The radiolabeled probes (200 bp) were generated by PCR using (γ-32P) dCTP 0.1 mM (3000 Ci. mmol-1 PerkinElmer). The primes used were: BCY1: Fw: 5′ CGAACAGGACACTCACCAGC 3′, Rv: 5′ GGTATCCAGTGCATCGGCAAG 3′; ENO2: Fw +1241 5’ GAGAATCGAAGAAGAATTGGGTG 3’; Rv +1469 5’ GCAATAG ACAGCAC GAGTCTTTG 3’; Rv +1336 5’ GCCGAGAGTCTTTTGGACTTTG 3’; TPK1: Fw: 5′ CCGAAG CAGCCACATGTCAC 3′; Rv: 5′ GTACTAACGACCTCGGGTGC 3′; TUB1: Fw: 5′ CAAGGG TTCTTGTTTACCCATTC 3′; Rv: 5′ GGATAAGACTGGAGAATA TGAAAC 3′. After washing, the membranes were subjected to digital image analysis (Bio-Imaging Analyzer Bas-1800II and Image J, National Institute of Health). The intensity at time 0 (before adding the drug) was defined as 100%, and the intensities at the other time points were calculated relative to time 0. The mRNA half-lives were determined by the best linear fits of the mRNA band intensities normalized to 28S and 18S rRNA loading control using the equation Rate = ln (2)/(t1/2).

### Microscopy

Yeast cell cultures were grown in SC + 2% glucose medium up to OD600 0.8–1 at 25 °C and subjected or not (control) to heat shock at 37 °C and treated or not with cyclohemide. Cells used for fluorescence microscopy were previously fixed with 7.4% formaldehyde and washed with P-bodies. All images were acquired using a spectral confocal microscope FluoView1000 and acquisition software FV10-ASW 2.0 (Olympus Co., Japan), employing a 60X UPLSAPO, NA 1.35 oil immersion objective. For each data acquisition, a z-stack of optical sections was acquired, in 1 µm or 0.5 µm steps, from the base of the cells on the coverslip to a total thickness of 9 µm. To avoid crosstalk, images were recorded in a sequential order. GFP and CFP were excited using the 488 nm and 458 lines of an argon laser and the emitted fluorescence were detected in the 500– 530 nm and 465–480 nm range, respectively. For samples with mCherry or RFP, 543 nm He-Ne was employed, and the emitted fluorescence was detected in the 580–680 nm or 555–655 nm range, respectively. ImageJ (http://rsbweb.nih.gov/ij/) was used to obtain equal contrast and adjust all images. As indicated in each figure, representative cells from independent cultures are shown. Cycloheximide treatments (100 µg/ml) were performed with minor modifications as previously described (Teixeira et al., 2005). For clarity the images shown are single Z-sections.

### Polysome profiling

Yeast cell cultures were grown in SC medium up to OD600 0.8–1 at 25°C and subjected or not (control) to heat shock (60 min at 37°C). 10 μg/ml cycloheximide was added and cells were lysed with glass beads. 1.5-2 A260 units of pre-cleared lysates were loaded onto 15–50% linear sucrose gradients. The gradients were centrifugated 1h 20 min at 50.000 rpm using a P55ST2 rotor (Hitachi 100NX), and then were fractionated from the bottom. 200 µl of each fraction were collected in a 96 well microplate absorption at 260 nm was measured to quantify RNA in a Microplate Reader DR-200Bs X. The tubes were snap-frozen in liquid N2 and stored at -80°C. RNA was extracted by isopropanol precipitation followed by the hot phenol method. Finally, mRNA levels were analysed by quantitative RT–PCR.

### Glycogen accumulation assay

Mid log cells were exposed to the scheme of temperature shifts. Samples were harvested at t=0 and after the first (i) and second (ii) incubation at 25°C. Cells collected were resuspended in a 0.2% iodine/0.4% potassium iodide solution or dilutions (1/2 and 1/4), incubated 3 min and then spotted onto YPD plates. The darker the color, the higher the amount of glycogen accumulated, providing an indirect measurement of less PKA activity

### Formaldehyde Cross-Linking

For cross-linking, yeast cells were pelleted and resuspended formaldehyde solution. Cells were incubated with mild agitation for 7 min at RT and then pelleted at 7000 g at RT for 3 min, resulting in 10 minutes exposure to formaldehyde. The supernatant was removed, and the reaction was quenched with 0.5 ml ice-cold 1.25M glycine/PBS. Cells were transferred to a smaller tube, spun, washed once in 1.25M glycine/PBS and lysed in 1ml RIPA buffer (50mM Tris HCl, pH 8.0, 150mM sodium chloride, 1%NP40, 0.5% sodium deoxycholate, 0.1% SDS, 1mM EDTA, protease inhibitors. Lysates were spun for 30 minutes at 20000 g at 4◦C to remove insoluble debris. The supernatant was either used directly or stored at −80◦C. Cell extracts were heated at 100°C or 65°C and analysed by Western blot using anti-Bcy1. The amounts of Tpk1 and Bcy1 were analyzed by SDS-PAGE following immunoblot with anti-Tpk1 or anti-Bcy1 (left panel). The quantification of Tpk1 and Bcy1 in the holoenzymes was performed using Image J.

### Author contributions

Luciana Cañonero: Investigation, Formal Results Analysis and validation, Constanza Pautasso: Investigation, Formal Results Analysis, Validation. Fiorella Galello: Review & Editing. Lorena Sigaut and Lia Pietrasanta: confocal microscopy assays. Javier Arroyo: Writing-Review, Analysis and discussion. Mariana Bermudez Moretti: Writing-Reviewing. Paula Portela: Writing-Review, Visualization, Analysis, and discussion. Silvia Rossi: Methodology, Conceptualization, Supervision, Writing - Original Draft, Writing - Review & Editing, Resources, Formal Analysis, Conceptualization, Methodology, Funding Acquisition.

## Acknowledgements

This work was supported by grants from the Agencia Nacional de Promoción Científica y Tecnológica (PICT 2014-2937, PICT 2017-02240 and PICT 2018-0378), from the University of Buenos Aires (UBA 2016–2018, 20020150100035BA). The work at the Javier Arroyo lab is supported by grants BIO2016-79289-P and PID2019-105223GB-I00 (Ministerio de Ciencia e Innovación, MICINN, Spain) and S2017/ BMD3691-InGEMICS (Comunidad de Madrid, Spain). Luciana Cañonero was awarded with PhD fellowships from CONICET. Constanza Pautasso was awarded with Pos-doctoral fellowships from CONICET. We thank Dr. Roy Parker from the University of Colorado Boulder for the generous gift of the pRP1661 plasmid.

## FIGURE LEGENDS

**Supplementary Figure 1.**
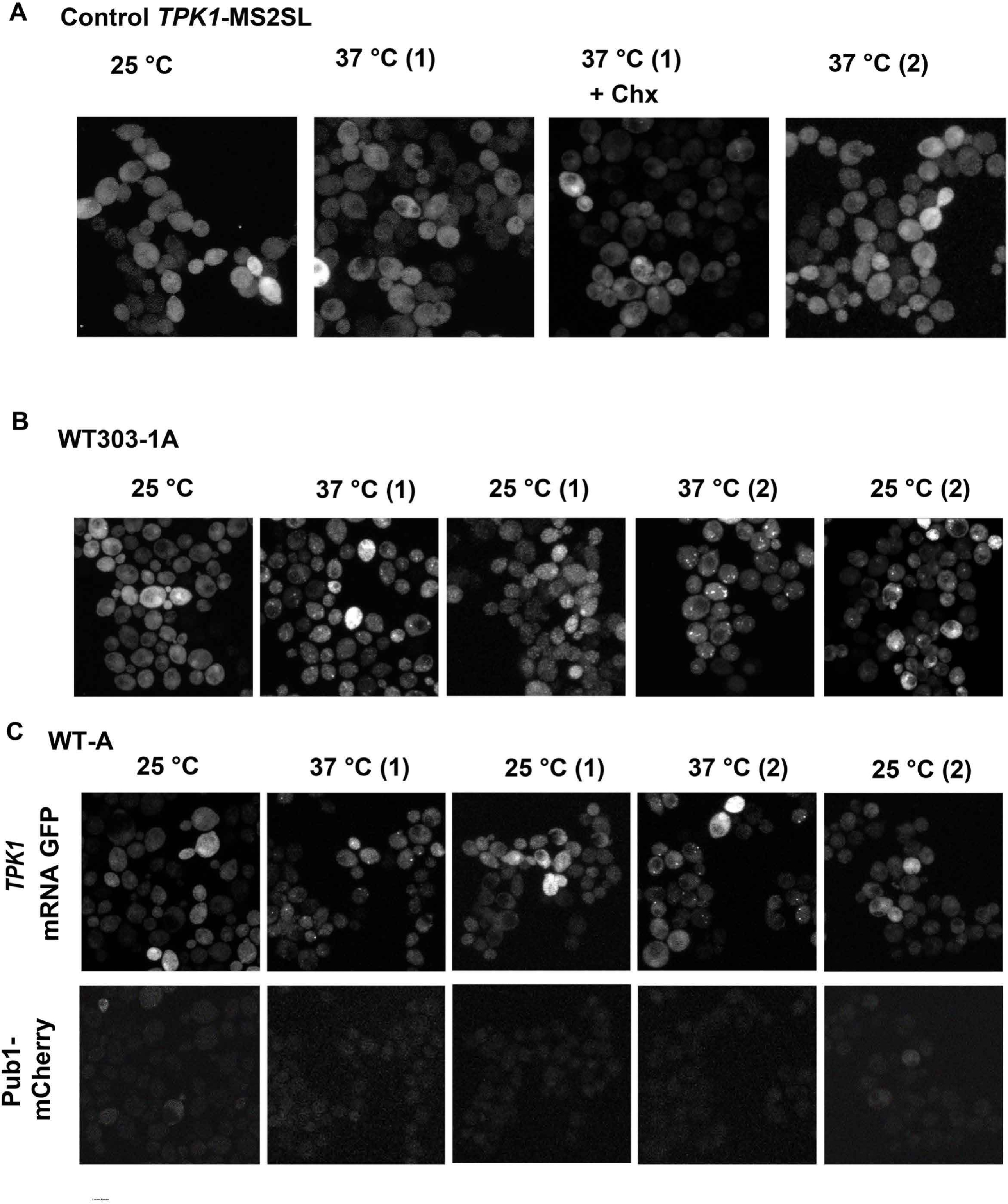
A) Control cells expressing only the pMS2-CP-GFPx3 but lacking MS2SL on *TPK1* (control *TPK1*-MS2SL) were incubated at 37°C for 60 minutes in the presence or not of cycloheximide (Chx). B and C) WT 303-1A and WT-A cells co-expressing *TPK1* mRNA-MS2SL and pMS2-CP-GFPx3 and neither Dcp2-CFP nor Pub1-mCherry were treated following the scheme of temperature shifts presented in Fig. 2A.

**Supplementary Figure 2.**
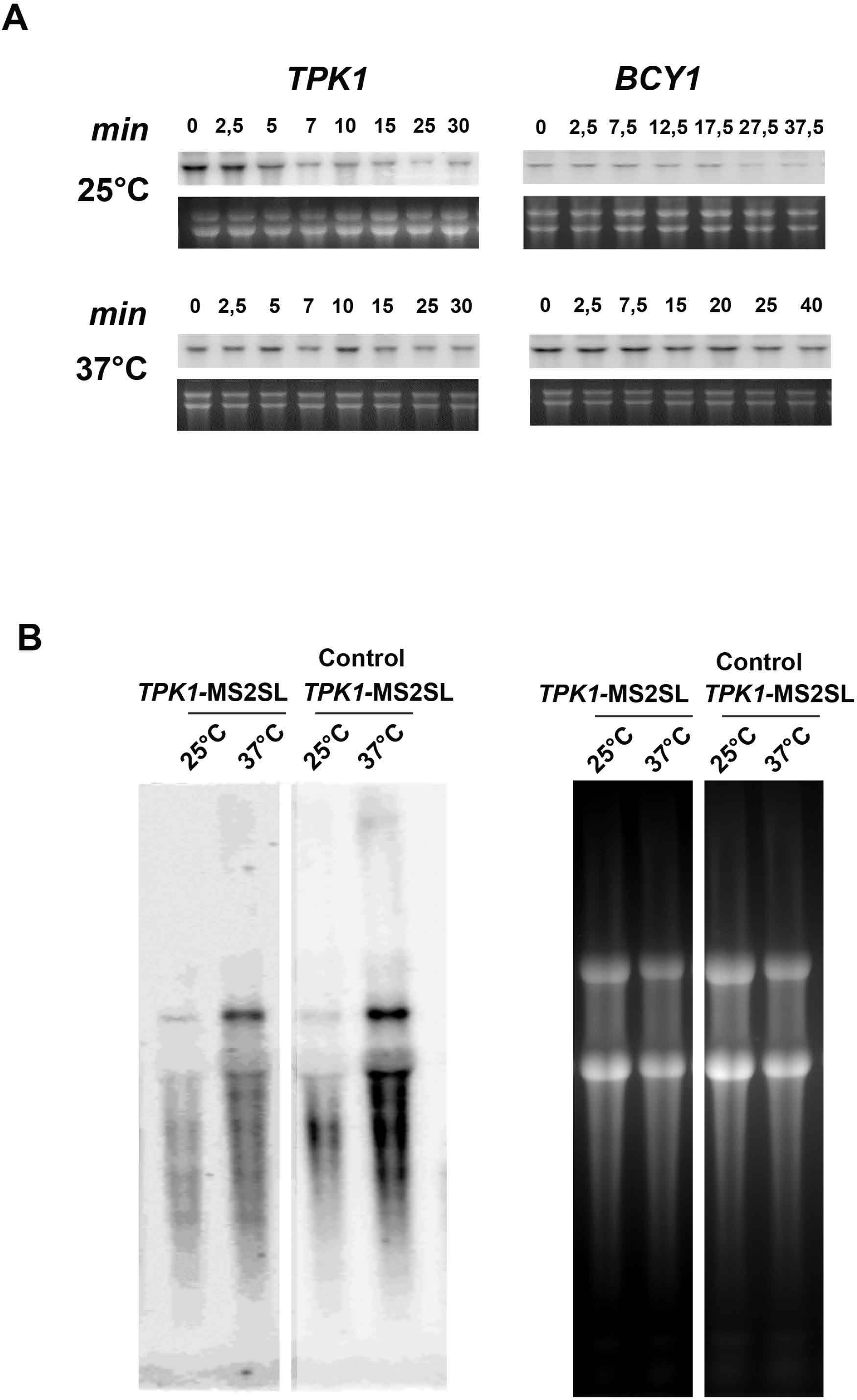
A) Cells were incubated or not at 37°C for 60 min and samples were harvested at the indicated time points following transcriptional arrest by 1,10-phenanthroline. *TPK1* mRNA levels at the indicated time points after the addition of the drug was analised by Northern blot. B) Cells co-expressing *TPK1* mRNA-M2SL and pMS2-CP-GFP (*TPK1*-M2SL) or expressing only *TPK1* mRNA-MS2SL (control *TPK1*-M2SL) were incubated at 37°C for 60 min and RNA integrity was determined by monitoring *TPK1* mRNA-MS2SL using a specific radiolabeled probe corresponding to the MS2SL sites.

**Supplementary Figure 3.**
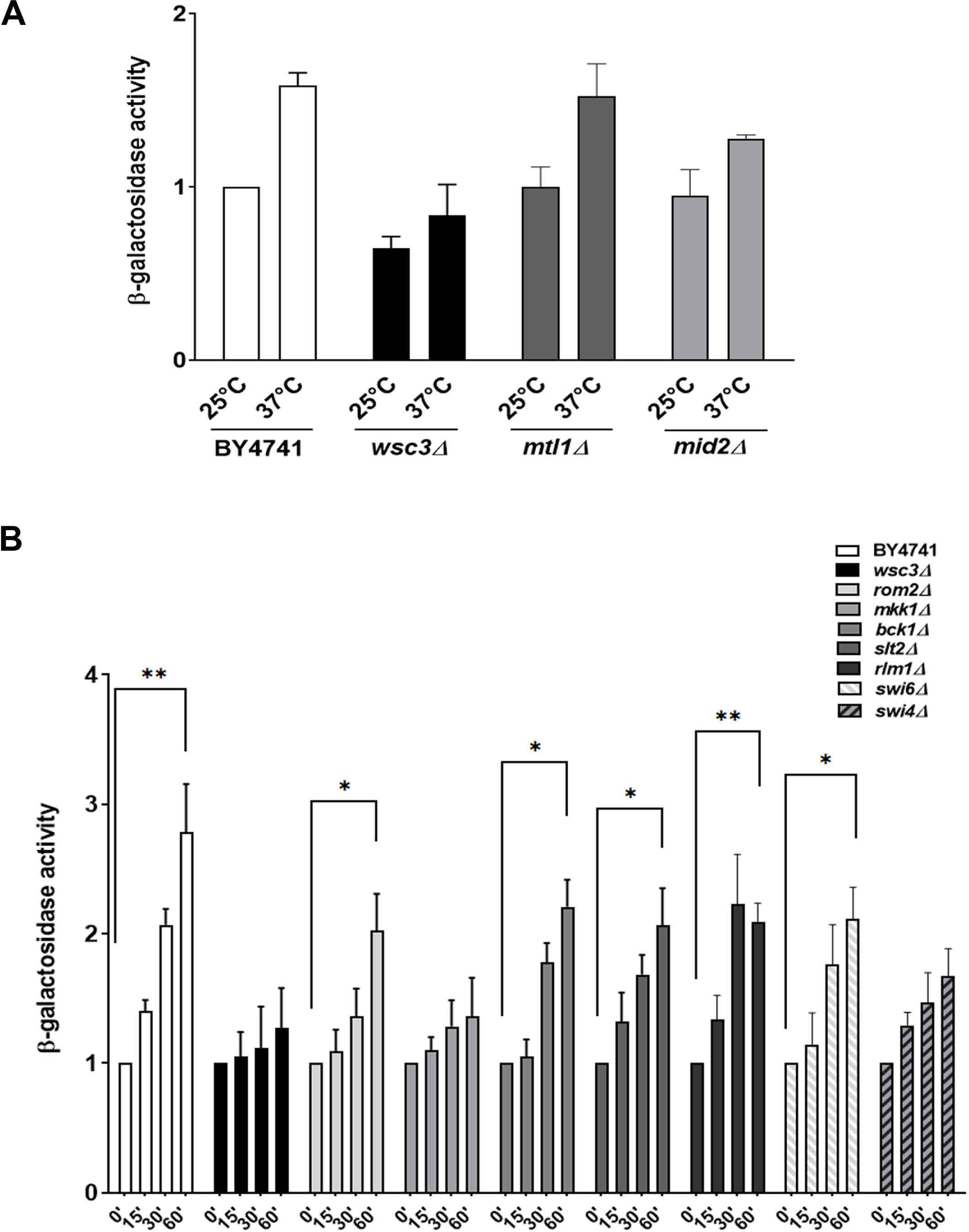
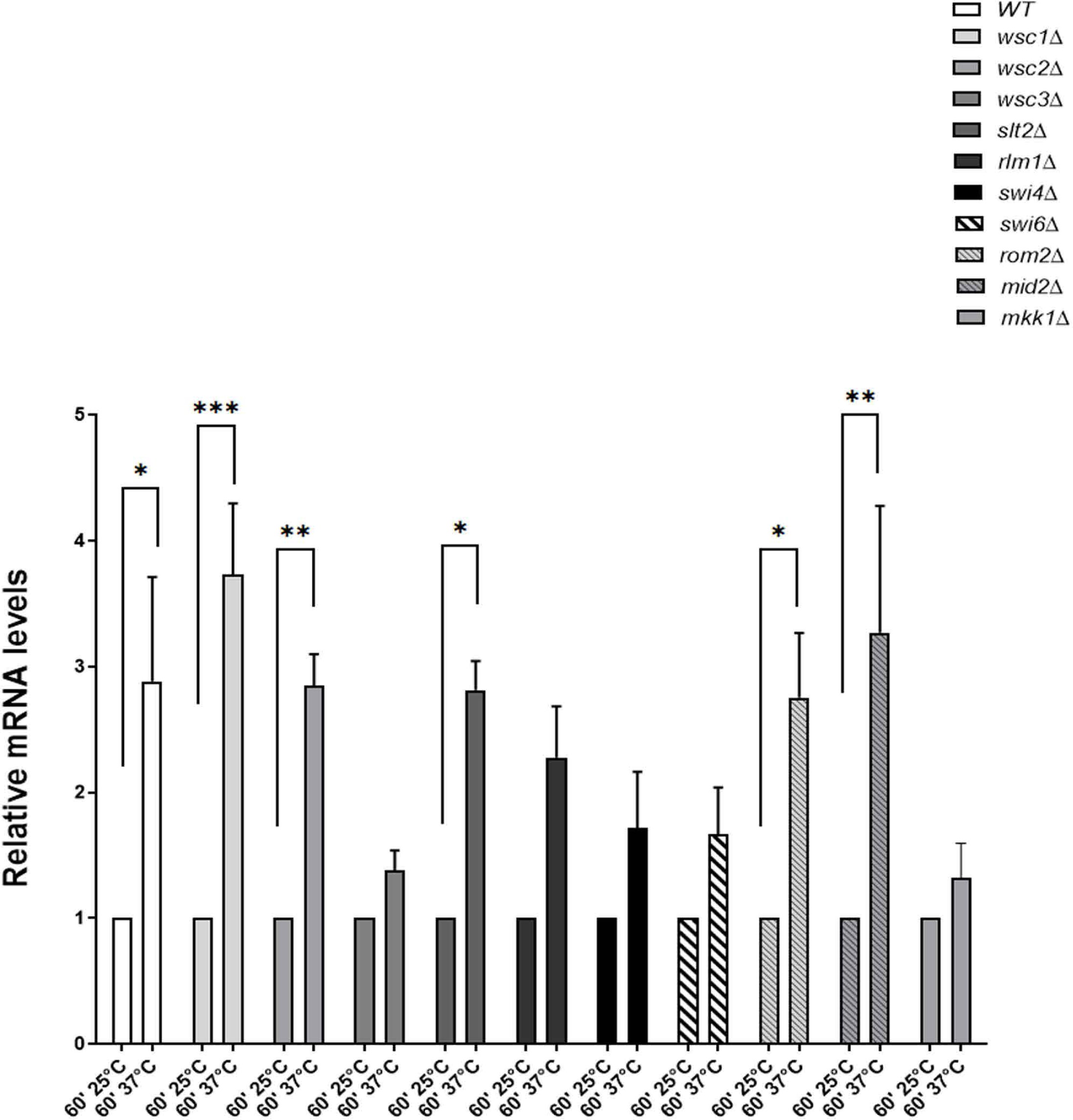
A and B) β-Galactosidase activity was determined in WT (BY4741), *wsc3Δ, mid2Δ* and *mtl1Δ*, *rom2Δ, bck1*Δ, *slt2*Δ, *rlm1*Δ, *swi4*Δ and *swi6*Δ strains carrying *TPK*-lacZ fusion gene. The results are expressed in Miller Units and as the mean ± s.e.m. of replicate samples (n= 4) from independent assays. C) *TPK1* mRNA levels of mutant strains used in A and B were determined by RT-qPCR, normalized to *TUB1* mRNA and expressed relative to 25°C for each strain. The values represent the mean ± s.e.m., obtained from n=3 biological replicates. Statistical analysis corresponds to two-way ANOVA, *p<0.05; **p<0.01; ***p<0.001.

**Supplementary Figure 4.**
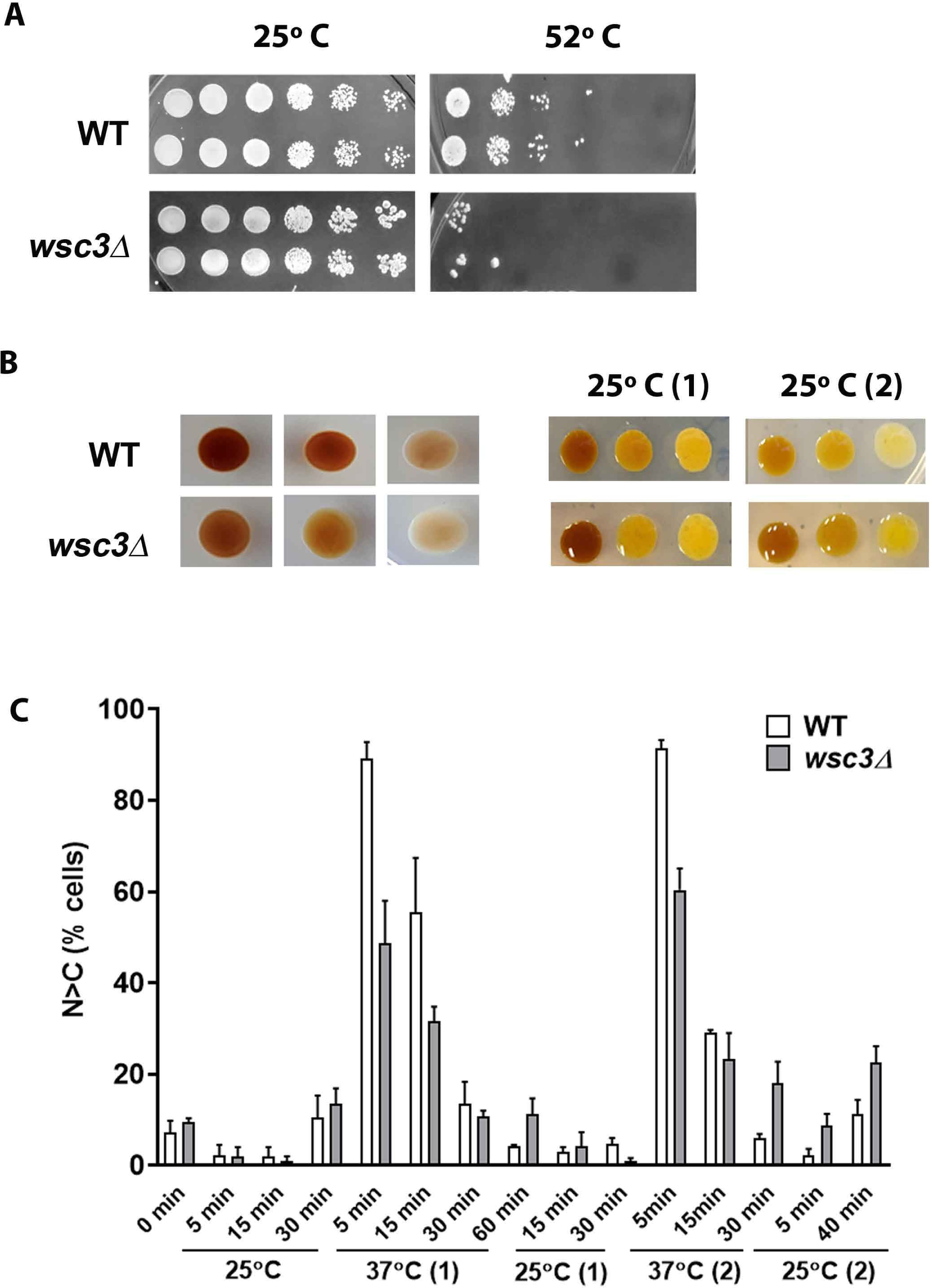
B) Cell viability assay of WT and *wsc3Δ* cells by spot assay. Cells were incubated at 52°C, 10 min and serial dilutions were spotted in YPD plates. B) Glycogen accumulation analysis, B) Bars graph depicts the percentage of cells showing nuclear Msn2-GFP localization in the wild-type strain and *wsc3Δ* treated at the indicated times and temperatures. The values represent the mean ± s.e.m., obtained from n=3.

**TABLE 1.**
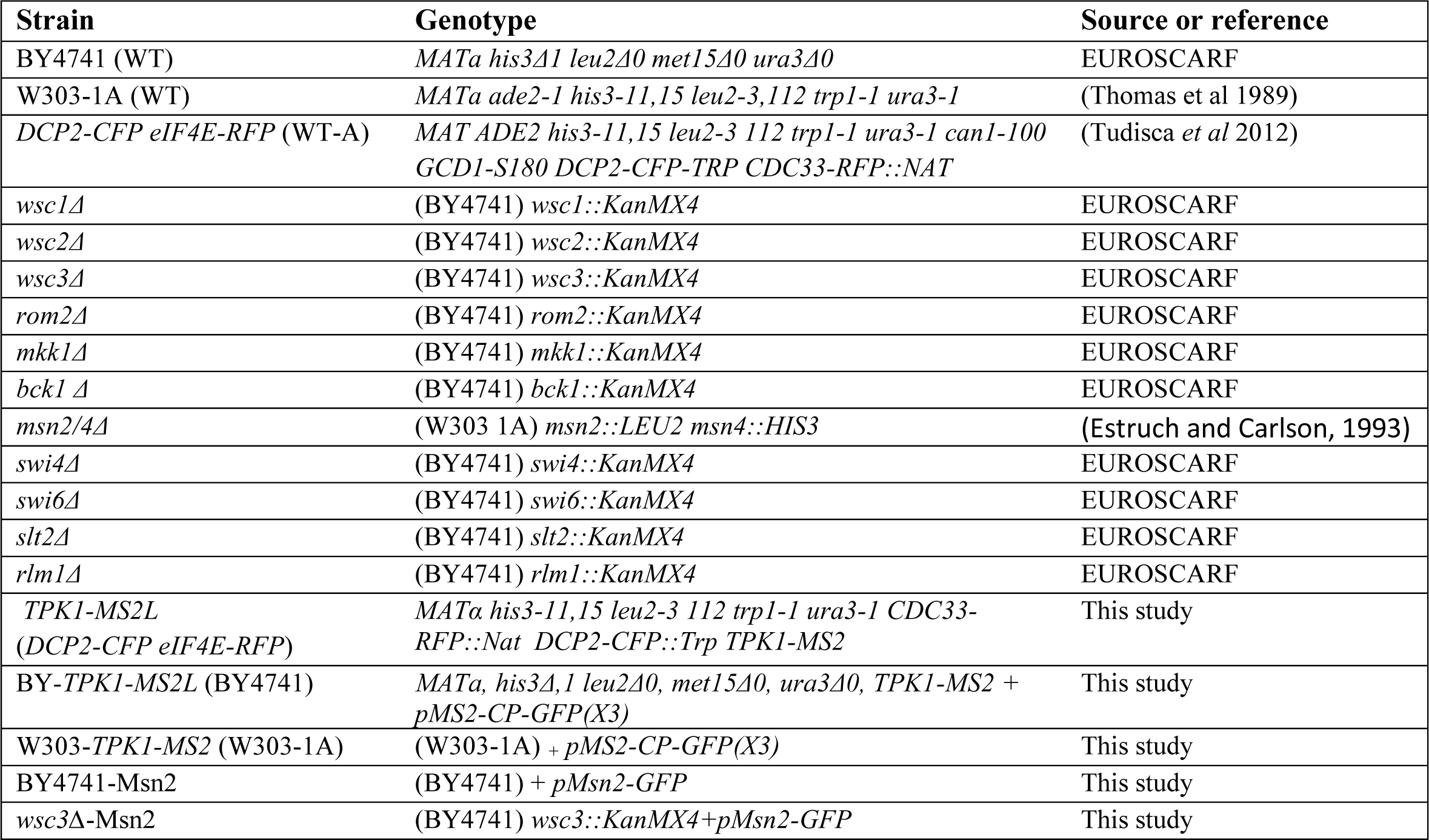
List of strains and nomenclature used in this study

**TABLE 2.**
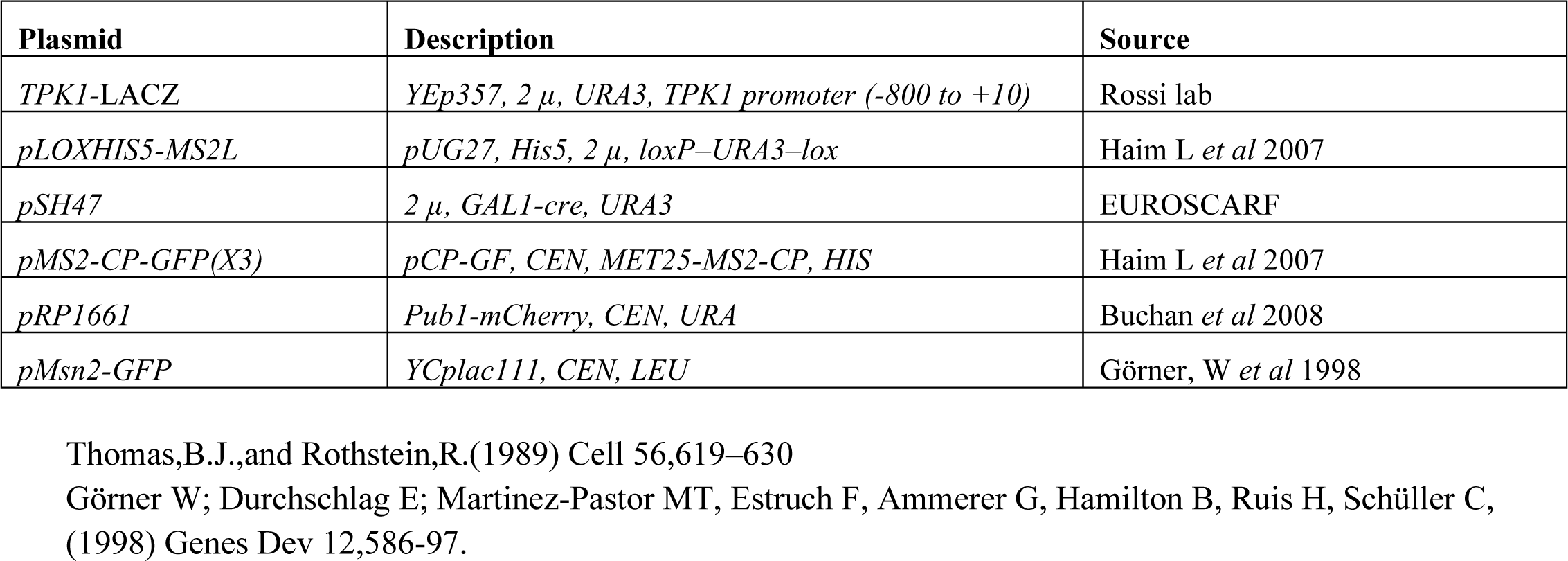
List of plasmids used in this study

## Checklist of key methodological and analytical information

This checklist is used to ensure good reporting standards and reproducibility in your paper (this checklist is compatible with the reporting standards recommended by the National Institutes of Health).

You must ensure that the following information is included in your manuscript. In general, this is best achieved by having specific subsections in the Materials and Methods section for reagents, animal models, statistics and data availability.

## Reagents

1. For cell lines, detail their source and state whether they were recently authenticated and tested for contamination. Confirm - in Materials & Methods: ■ Reported elsewhere (specify)/NA:
2. For antibodies, provide a citation, catalogue number and/or clone number and batch number. Provide details on antibody validation, either in Supplementary Information or by referencing an antibody validation profile (where possible). Give the dilutions used. Confirm – in Materials & Methods:■ Reported elsewhere (specify)/NA:

## Animal models

We recommend consulting the ARRIVE guidelines to ensure that other relevant aspects of animal studies are adequately reported.

1. Report species, strain, sex and age of animals. Confirm – in Materials & Methods:■ Reported elsewhere (specify)/NA:
2. Provide details on compliance with relevant ethical regulations including, where necessary, the identity of the committee(s) approving the experiments. Confirm – in Materials & Methods:■ Reported elsewhere (specify)/NA:

## Human subjects

1. Provide details on compliance with relevant ethical regulations and identify the committee(s) approving the study protocol.
2. Provide a statement confirming that informed consent was obtained from all subjects.
3. Where photographs of patients are included, provide a statement confirming that consent to publish was obtained.
4. For work involving human eggs or embryos, any financial recompense to donors must be declared.
5. Where the work reports new clinical trial data or includes a tumour marker prognostic study, appropriate guidelines for reporting must be followed (e.g. reporting the clinical trial registration number, submitting a CONSORT checklist, following REMARK reporting guidelines). Please contact the editorial office for further guidance if required. Confirm – in Materials & Methods:■ Reported elsewhere (specify)/NA:

## Data availability

For further details on our policies regarding data availability, please see <u>here</u>.

1. Include accession codes for deposited data. Confirm – in Materials & Methods:■ Reported elsewhere (specify)/NA:
2. Include the source of all software. For any custom software, include a statement of how it can be obtained. Confirm – in Materials & Methods:■ Reported elsewhere (specify)/NA:

## Methodology and statistics

The Materials and Methods section should provide information on all points listed below. Please read these carefully and confirm that your manuscript conforms to these standards.

1. State how the sample size (*n*) was defined to ensure adequate power to detect a pre-specified effect size.
2. Describe inclusion and exclusion criteria if samples or animals were excluded from the analysis. State whether the criteria were pre-established.
3. Describe any methods of randomisation used to determine how samples or animals were allocated to experimental groups and processed.
4. If the investigator was blinded to the group allocation during the experiment and/or when assessing the outcome, state the extent of blinding.
5. For data presented, statistical tests must be appropriate to the type of data. For example, do the data meet the assumptions of the tests (e.g. normal distribution)? Is there an estimate of variation within each group of data? Is the variance similar between the groups that are being statistically compared?

For small sample sizes (*n*<5), descriptive statistics are not appropriate, and instead individual data points should be plotted.

Confirm:■ or Reported elsewhere (specify):

## Figure legends

The following should be reported in every figure legend.

1. The exact sample size (*n*) for each experimental group or condition, given as a number, not a range.
2. A description of the sample collection that allows the reader to understand whether the samples represent technical or biological replicates (including how many animals, cultures, etc.).
3. A statement of how many times the experiment shown was replicated in the laboratory.
4. Definition of average values as median or mean; definition of error bars as s.d., s.e.m. or c.i. (please write as e.g. mean±s.e.m.). Error bars should reflect independent experiments and not technical replicates.
5. Statistical test results, e.g. *P* values.
6. Details of statistical method.
  - *t*-test, simple χ2 tests, Wilcoxon, Mann–Whitney tests and one-way and two-way ANOVA tests can be identified by name only in the figure legend. More complex tests should be described in the Materials and Methods.
  - Are tests one-tailed or two-tailed?
  - Are there adjustments for multiple comparisons?

Confirm:■ or Reported elsewhere (specify):

